# An integrated approach unravels a crucial structural property for the function of the insect steroidogenic Halloween protein Noppera-bo

**DOI:** 10.1101/781070

**Authors:** Kotaro Koiwai, Kazue Inaba, Kana Morohashi, Sora Enya, Reina Arai, Hirotatsu Kojima, Takayoshi Okabe, Yuuta Fujikawa, Hideshi Inoue, Ryunosuke Yoshino, Takatsugu Hirokawa, Koichiro Kato, Kaori Fukuzawa, Yuko Shimada-Niwa, Akira Nakamura, Fumiaki Yumoto, Toshiya Senda, Ryusuke Niwa

## Abstract

Ecdysteroids are the principal insect steroid hormones essential for insect development and physiology. In the last 18 years, several enzymes responsible for ecdysteroid biosynthesis, encoded by Halloween genes, have been identified and well characterized, both genetically and biochemically. However, none of these proteins have yet been characterized at the tertiary structure level. Here, we report an integrated *in silico*, *in vitro*, and *in vivo* analyses of the Halloween glutathione *S*-transferase (GST) protein, Noppera-bo (Nobo). We determine crystal structures of *Drosophila melanogaster* Nobo (DmNobo) complexed with glutathione and 17β-estradiol, a DmNobo inhibitor. 17β-estradiol almost fully occupied the putative ligand-binding pocket, and a prominent hydrogen bond formed between Asp113 of DmNobo and 17β-estradiol. Asp113 is essential for inhibiting DmNobo enzymatic activity by 17β-estradiol, as 17β-estradiol does not inhibit and physically interacts less with the Asp113Ala DmNobo point mutant. Asp113 is highly conserved among Nobo proteins, but not among other GSTs, implying that Asp113 is important for endogenous Nobo function. Indeed, a homozygous *nobo* allele possessing the Asp113Ala point mutation exhibits embryonic lethality with undifferentiated cuticle structure, a phenocopy of complete loss-of-function *nobo* homozygotes. These results suggest that the *nobo* family of GST proteins has acquired a unique amino acid residue, which seems to be essential for binding an endogenous sterol substrate to regulate ecdysteroid biosynthesis. This is the first study to reveal the structural characteristics of insect steroidogenic Halloween proteins. This study also provides basic insight into applied entomology for developing a new type of insecticides that specifically inhibit ecdysteroid biosynthesis.

**Significance Statement:** Insect molting and metamorphosis are drastic and dynamic biological processes and, therefore, have fascinated many scientists. Ecdysteroids represent one class of insect hormones that are indispensable for inducing molting and metamorphosis. It is well known that proteins responsible for catalyzing ecdysteroid biosynthesis reactions are encoded by “Halloween” genes, most of which have names of ghosts and phantoms. However, no studies have focused on the structural properties of these biosynthetic proteins. In this study, we addressed this unsolved issue and successfully unraveled a structural property that is crucial for the function of the fruit fly Halloween protein, Noppera-bo (a Japanese faceless ghost). This is the first study to reveal the structural characteristics of an insect steroidogenic Halloween protein.

## Introduction

Ecdysteroids play pivotal roles in regulating many aspects of development and physiology in arthropods, including insects (1, 2). Because ecdysteroids do not exist naturally in animals other than arthropods, it has been long thought that molecules involved in ecdysteroid biosynthesis, secretion, circulation and reception could be good targets for developing third-generation pesticides that specifically inhibit insect life cycles, with no adverse effects on other animals (3). Thus, the study of ecdysteroids has been important, not only in the basic biological sciences, but also in the field of applied agrobiology.

Ecdysteroids are biosynthesized from dietary sterols that are primarily obtained from food sources (1, 2). The formation of each biosynthetic intermediate going from dietary sterols to the biologically-active form of ecdysteroids, 20-hydroxyecdysone (20E), is catalyzed by a specific ecdysteroidogenic enzyme (2, 4). Since 2000, a series of these enzymes has been identified. These enzymes include Neverland (5, 6), Non-molting glossy/Shroud (7), Spook/CYP307A1 (8, 9), Spookier/CYP307A2 (9), CYP6T3 (10), Phantom/CYP306A1 (11, 12), Disembodied/CYP302A1 (13), Shadow/CYP315A1 (13), and Shade/CYP314A1 (14). A deficiency of genes encoding these enzymes results in developmental lethality. Particularly, in the fruit fly *Drosophila melanogaster*, complete loss-of-function mutants of *shroud*, *spook*, *phantom*, *disembodied*, *shade*, and *shadow*, which are often classified as Halloween mutants, commonly result in embryonic lethality with the loss of differentiated cuticle structures (15). To date, the functions of these enzymes have been characterized genetically and some of them have also been analyzed biochemically (2, 16). However, none of these enzymes have yet been characterized at the tertiary structure level.

Here, we report the first crystal structure of an ecdysteroidogenic regulator encoded by the Halloween gene, *noppera-bo* (*nobo*) (17–19). *nobo* encodes a member of the epsilon class of cytosolic glutathione *S*-transferases (GST, EC 2.5.1.18; hereafter GSTEs) (20). In general, GSTs catalyze various reactions with an activated glutathione (GSH) molecule in the following 3 ways: GSH conjugation to a substrate, reduction of a substrate using GSH, and isomerization (21). Data from previous studies have demonstrated that *nobo* is specifically expressed in ecdysteroidogenic tissues, including the prothoracic gland and the adult ovary (17–19). In addition, loss-of-*nobo*-function mutations in *D. melanogaster* and *Bombyx mori* result in developmental lethality, which are well rescued by administering 20E (17–19). Consistent with the requirement of GSH for GST function, a defect in glutathione biosynthesis in *D. melanogaster* also leads to larval lethality, in part due to ecdysone deficiency (22). These data clearly indicate that the *nobo* family of GSTs is essential for ecdysteroid biosynthesis. However, besides GSH, an endogenous ligand and a catalytic reaction driven by Nobo have not been elucidated.

In this study, we utilized the vertebrate female sex hormone 17β-estradiol (EST, Fig. 1*A*) as a molecular probe to gain insight into Nobo ligand recognition, based on our previous finding that EST inhibits the GSH-conjugation activity of *D. melanogaster* Nobo (DmNobo; also known as DmGSTE14) (23). We therefore considered the complex of DmNobo and EST to be an ideal target for elucidating a 3-dimentional structure of an ecdysteroidogenic Halloween protein and characterizing the interaction between DmNobo and its potent inhibitor. Moreover, we used an integrated, combined approach based on quantum chemical calculations, molecular dynamics (MD) simulations, biochemical and biophysical analyses, and molecular genetics. Consequently, we identified one DmNobo amino acid residue that is strongly conserved only in the *nobo* family of GSTs, which is crucial for DmNobo inhibition by EST and for the normal *in vivo* function of DmNobo during *D. melanogaster* embryogenesis.

**Figure 1.**
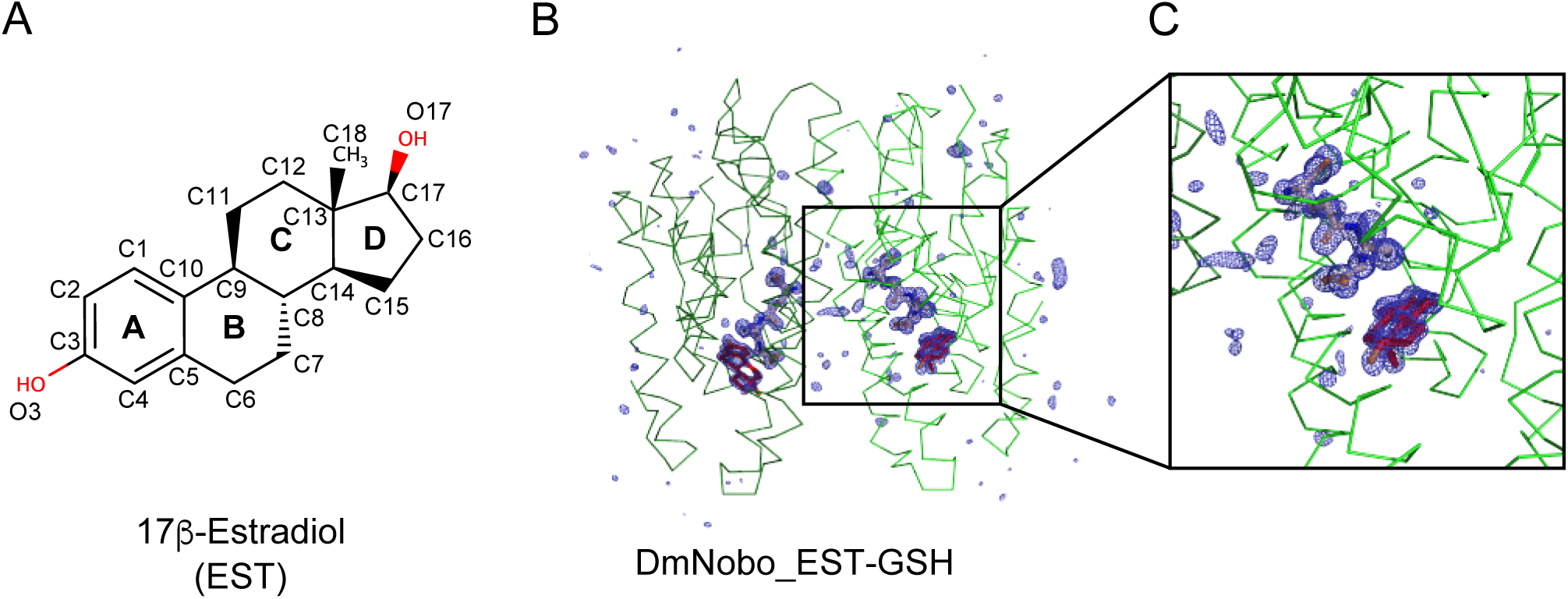
Crystal structures of the *Drosophila melanogaster* Noppera-bo protein. (A) Chemical structure of 17β-estradiol (EST). The atoms of the steroid nucleus are indicated. Rings A, B, C, and D are also shown. (B) Simulated annealing-omit map for GSH and EST in the DmNobo_EST-GSH complex. A m*F*o-D*F*c map (blue) (4.0σ) within 5.0 Å from the protein atoms is shown. Carbon atoms of DmNobo, GSH, and EST are colored green, wheat, and red, respectively. Oxygen and nitrogen atoms are colored green and blue, respectively. (C) An enlarged view of (B) around the EST and GSH ligands

## Results

### Crystal structure of DmNobo

The crystal structure of the apo-form of DmNobo (DmNobo_Apo) was determined at 1.50-Å resolution by the molecular replacement method (SI *Appendix*, Fig. S1*A*, Table S1). DmNobo forms a polypeptide homodimer with a canonical GST fold, which has a well-conserved GSH-binding site (G-site) and a hydrophobic substrate-binding pocket (H-site) adjacent to the G-site (21, 24). The crystal structures of the DmNobo_GSH, DmNobo_EST, and DmNobo_EST-GSH complexes were also determined at resolutions of 1.75 Å, 1.58 Å, and 1.55 Å, respectively (Fig. 1*B*, SI *Appendix*, Fig. S1*B*, Table S1). The crystal structures of the DmNobo_EST and DmNobo_EST-GSH complexes reproducibly showed clear electron densities for EST. GSH and EST binding did not affect the overall structure of DmNobo (SI *Appendix*, Fig. S1*C*); the root-mean-square deviation (RMSD) values for each pair among the four crystal structures were comparable with respect to the estimated coordinate errors (SI *Appendix*, Table S2).

GSH, a common substrate of GSTs (21, 24), was found in the G-site of DmNobo. Crystallographic analysis revealed that the position and conformation of GSH in DmNobo were essentially identical to those in other GSTEs (25–27). GSH is recognized by an intensive hydrogen bond network with Gln43, His55, Val57, Pro58, Asp69, Ser70, His71, and Ser107 in the G-site (SI *Appendix*, Fig. S2). The hydrogen bond interactions are similar to those found in other GSTE proteins (25–27). Moreover, these residues are well conserved among not only GSTEs but also the delta and theta classes of GSTs (hereafter GSTD proteins and GSTT proteins, respectively), which are closely related to GSTEs (SI *Appendix*, Fig. S3*A* and S3*B*) (20). Therefore, we conclude that the interaction between the G-site and GSH cannot account for the unique functional property of DmNobo, as compared to other GSTD/E/T proteins.

### Molecular mechanism of EST recognition by DmNobo

EST was bound in the H-site, which has a hydrophobic character. The electron-density map clearly showed that the compound in the H-site was the intact EST molecule. The EST molecule had no chemical modifications, including reduction and *S*-glutathionylation. The H-site, of which volume is approximately 365 Å^3^, was mostly filled with the EST molecule, which has a volume of approximately 350 Å^3^, and no space was available to accommodate another compound in the H-site (SI *Appendix*, Fig. S4).

Of the 16 amino acid residues lining the H-site, Arg13, Ser14, Gln43, Ser118, Arg122, and Met212 do not have direct contacts with EST (SI *Appendix*, Table S3). The D-ring of EST is situated near the entrance of the H-site and exposed to the solvent. Only a few interactions are observed between the D-ring of EST and DmNobo (Fig. 2*A*, SI *Appendix*, Table S3). In contrast, the A-ring of EST is located deep inside of the H-site and makes intensive interactions with H-site residues (Fig. 2*A*, SI *Appendix*, Table S3). Ser14, Pro15, Leu38, Phe39, and Phe110 form hydrophobic interactions with the A-ring of EST and are completely conserved among the Nobo proteins (Fig. 3*A*, Fig. 3*B*, Fig. 3*C*, SI *Appendix*, Table S3). The conservation of these residues among DmGSTD/E/T proteins are less than those among Nobo proteins (Fig. 3*D*, Fig. 3*E*, Fig. 3*F*, SI *Appendix*, Table S3). While Ser114, Met117, Ser118, Val121, Thr172, and Leu208, which form hydrophobic interactions with EST, are not conserved completely in the Nobo proteins, they show higher conservation than that found among DmGSTD/E/T proteins. These results suggest that the three-dimensional structure of the H-site, particularly near the A-ring of EST, is conserved in Nobo proteins and has different characteristics from DmGSTD/E/T proteins.

**Figure 2.**
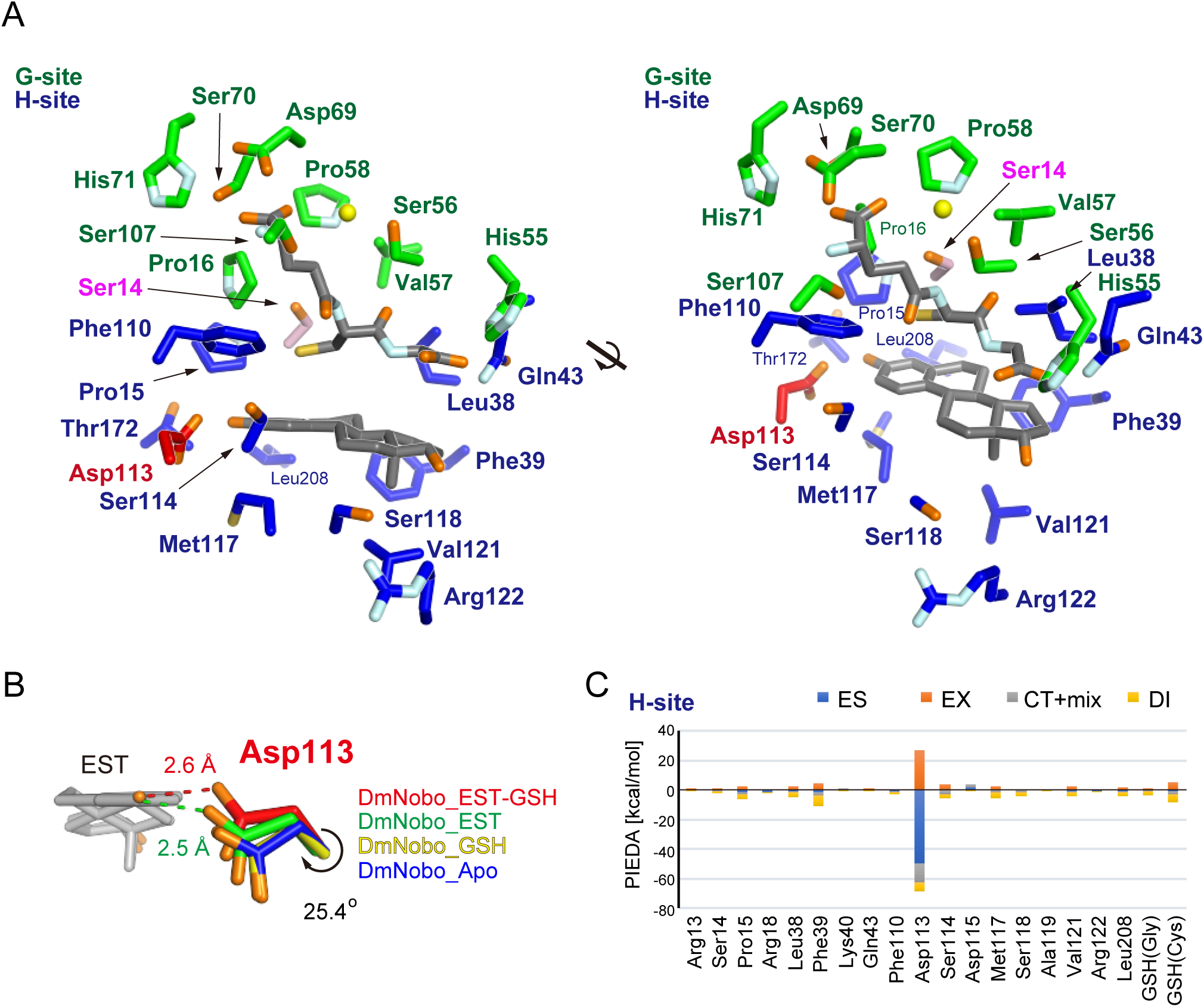
Asp113 in the H-site interacts with 17β-estradiol. (A) GSH- and EST-interacting residues. Carbon atoms of the G- and H-sites are colored in green and blue, respectively. Common residues of the G- and H-sites (Ser14, Pro15, Leu38, Gln43, and Phe110) are assigned as those of the H-site in this figure. Carbon atoms in Ser14, Asp113, and ligands (GSH and EST) are colored in pink, red, and gray, respectively. A water molecule interacting with each ligand is represented with a yellow sphere. (B) Conformational change of Asp113 upon ligand binding. Carbon atoms in DmNobo_Apo, DmNobo_GSH, DmNobo_EST, and DmNobo_EST-GSH are shown in blue, yellow, green, and red, respectively. A hydrogen bond between the O3 atom of EST and Oδ in Asp113 is indicated by a dashed line. The difference in the χ1 torsion angle of Asp113 between DmNobo_GSH and DmNobo_EST-GSH was 25.4°. (C) Interaction energies between EST and other atoms in the DmNobo_EST-GSH complex. The interaction energies were calculated from the PIEDA analysis, based on the FMO calculation. ES, EX, CT+mix, and DI indicate the electrostatic energy, exchange repulsion energy, charge transfer energy and higher order mixed term, and dispersion energy, respectively. Residues within a distance of twice the van der Waals radii from the EST atoms are shown. Numerical data for (C) are available in the SI *Appendix*, Table S4.

**Figure 3.**
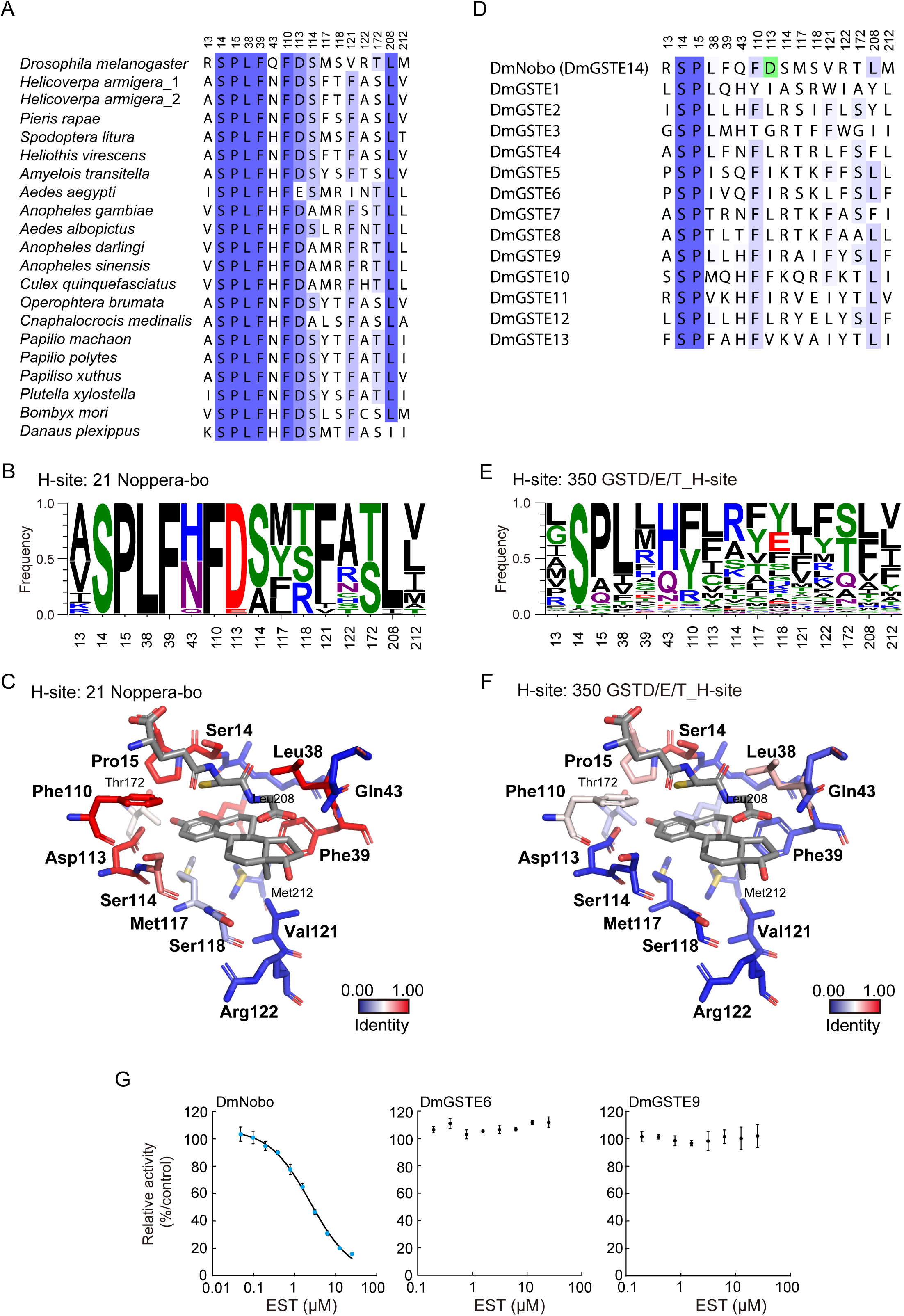
Consensus amino acid residues in the H-sites of Nobo orthologues. (A) Amino acid-sequence alignment of the H-site residues of 21 Nobo orthologues. These sequences were aligned using COBALT and manually edited, based on the crystal structure of DmNobo. The accession numbers of Helicoverpa armigera_1 and _2 are XP_021192638.1 and A0A2W1BRE1, respectively. (B) Frequencies of amino acid residues forming the H-sites of 21 Nobo. The frequencies were calculated using LOGO. (C) Conservation ratios of H-site residues among Nobo proteins are mapped to the tertiary structure of DmNobo. (D) Amino acid-sequence alignment of the H-site residues of DmGSTE. Asp113 of DmNobo is colored in green. (E) Frequencies of amino acid residues forming the H-sites of GSTD/E/T proteins. The frequencies were calculated using LOGO. (F) Conservation ratios of H-site residues among GSTD/E/T proteins including Nobo proteins (SI Appendix, Fig. S3A, Table S2) are mapped to the tertiary structure of DmNobo. (G) EST-dependent inhibition of the GSH-conjugation activities of DmNobo, DmGSTE6, and DmGSTE9. 3,4-DNADCF was used as an artificial fluorescent substrate. Each relative activity is defined as the ratio of activity, when compared to the respective proteins without EST. The error bars indicate the standard errors (SEM) from triplicate assays.

While the H-site has an overall hydrophobic character, there is one charged residue, Asp113, in the H-site. Asp113, which is nearly completely conserved in the Nobo proteins (see below), is located at the innermost region of the H-site and forms a hydrogen bond with a hydroxyl group on the C3 atom of EST. EST binding induces a rotation of the χ1 angle of Asp113 by 25.4°, and Oδ of Asp113 forms a hydrogen bond with O3 of EST (Fig. 2*B*). The carboxyl group of Asp appears to be ionized, as expected considering that its isoelectric point is approximately 3.6 and that crystallization in solution was achieved at a pH of 6.4. This is the only hydrogen bond found between EST and DmNobo and seems to be critical for EST binding.

To evaluate the contribution of the hydrogen bond to the interaction with EST, total interaction energies between EST fragments and DmNobo amino acid residues were calculated using the fragment molecular orbital (FMO) method, which can evaluate the inter-fragment interaction energy (IFIE) based on the quantum chemistry (28, 29). The FMO calculation classifies the IFIE into 4 energy categories, namely the electrostatic energy (ES), exchange-repulsion energy (EX), charge-transfer energy and higher-order mixed term (CT+mix), and dispersion energy (DI). The FMO calculation estimated that the ES represented approximately half of the total IFIE (−41.4 kcal/mol versus −82.4 kcal/mol; Fig. 2*C* and SI *Appendix*, Table S4). The crystal structure suggested that the ES arises from the hydrogen bond between Oδ of Asp113 and O3 of EST (SI *Appendix*, Table S4). These results suggested that Asp113 plays a critical role in interacting with EST.

### Asp113 in DmNobo is essential for EST binding

The importance of the Asp113-EST hydrogen bond for EST binding was biochemically examined with a recombinant mutated DmNobo protein carrying Asp113Ala amino acid substitution (DmNobo[Asp113Ala]). DmNobo[Asp113Ala] lacks the sidechain carboxyl group at position 113 and therefore cannot form a hydrogen bond with EST. The crystal structure of the DmNobo[Asp113Ala] did not show significant structural differences compared with the wild-type DmNobo (DmNobo[WT]) protein (SI *Appendix*, Fig. S5*A*, Fig. S5*B*).

We first examined the enzymatic activities of DmNobo[WT] and DmNobo[Asp113Ala] using an *in vitro* enzymatic assay system with the fluorogenic substrate 3,4-DNADCF (23). In this assay system, GSTs catalyze GSH conjugation to the non-fluorescent molecule, 3,4-DNADCF, giving rise to highly fluorescent product, 4-GS-3-NADCF. In the absence of EST, both DmNobo[WT] and DmNobo[Asp113Ala] showed GSH-conjugation activity (Fig. 4*C*). In the presence of EST, as expected from the EST-binding to the H-site, the enzymatic activity of DmNobo[WT] was inhibited with an IC_50_ value of approximately 2.3 μM (Fig. 4*A*, Fig. 4*C*). In contrast, the enzymatic activity of DmNobo[Asp113Ala] was not inhibited by EST, even at a concentration of 25 μM (Fig. 4*A*, Fig. 4*C*).

**Figure 4.**
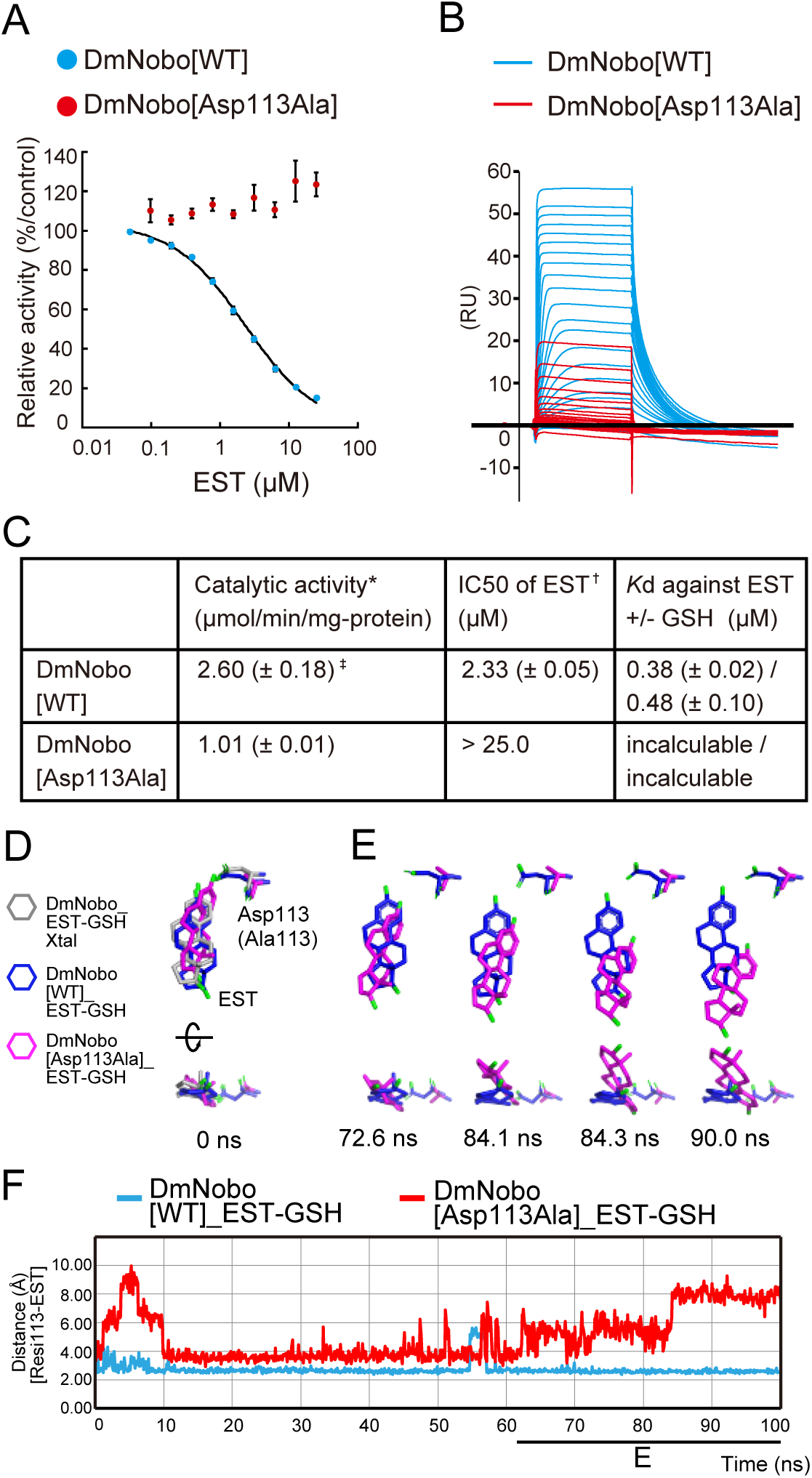
Asp113 is essential for DmNobo binding to EST. (A) EST-dependent inhibition of the GSH-conjugation activity of DmNobo[WT] (cyan) and DmNobo[Asp113Ala] (red). 3,4-DNADCF was used as an artificial fluorescent substrate. In each case, the relative activity is defined as the ratio of activity, when compared to DmNobo[WT] without EST. The error bars indicate the standard errors (SEM) from triplicate assays. (B) Sensorgrams of surface plasmon-resonance analysis of DmNobo proteins with EST. DmNobo[WT] or DmNobo[Asp113Ala] was immobilized to a sensor chip, and solutions containing a series of EST concentrations were applied in presence of 1 mM GSH. (C) Kinetic parameters of DmNobo proteins. Catalytic activity (*) and IC_50_ of EST (†) indicate 3,4-DNADCF-specific GSH-conjugation activity and the IC_50_ of EST against 3,4-DNADCF-specific GSH-conjugation activity, respectively. Values in parentheses indicate standard errors from triplicate assays (‡). (D–F) *In silico* evaluation of the contribution of Asp113 to the interaction between DmNobo and EST. MD simulations of the DmNobo[WT] or DmNobo[Asp113Ala] complex with EST and GSH in a TIP3P-water model were carried out at 300 K for 100 ns. These simulations were performed in triplicate. (D) MD models at 0 ns of DmNobo with EST and GSH (blue), DmNobo[Aps113Ala] with EST and GSH (magenta), and the crystal structure of DmNobo_EST-GSH (EST-GSH_Xtal, gray). The upper models are shown from above the EST ligand, and the lower models are rotated 90° from the upper models. Hydrogen atoms are not shown. (E) MD models of DmNobo[WT]_EST-GSH and DmNobo[Asp113Ala]_EST-GSH from 72.6 ns to 90.0 ns (F) Distance between Oδ of Asp113 of DmNobo[WT] or Cβ of DmNobo[Asp113Ala] and the O3 atom of EST at each frame.

We next measured the dissociation constant (*K*d) values between DmNobo and EST by performing surface plasmon-resonance (SPR) analysis. The *K*d values between DmNobo[WT] and EST in the presence or absence of GSH were 0.38 ± 0.02 µM and 0.48 ± 0.10 µM, respectively (Fig. 4*B*, Fig. 4*C*). In contrast, it was barely possible to determine the *K*d value between DmNobo[Asp113Ala] and EST due to a weak interaction (Fig. 4*B*, Fig. 4*C*), which was consistent with crystal structure analysis (SI *Appendix*, Fig. S5*C*). These results suggest that Asp113 is critical for interaction with EST.

We also employed MD simulations to confirm the contribution of Asp113 to the interaction with EST using DmNobo[WT] and DmNobo[Asp113Ala] as models. In these MD simulations, the initial structures of EST and the DmNobo proteins were defined based on data acquired from our crystallographic analyses (Fig. 4*D*). While simulating DmNobo[WT] for 100 nano seconds (ns), we found that the distance between Oδ of Asp113 and the hydroxyl group of EST was relatively constant (Fig. 4*E* and 4*F*, Movie 1, Movie 2). However, when simulating DmNobo[Asp113Ala], the distance between Ala113 and the hydroxyl group of EST increased over time, and EST moved from the initial position (Fig. 4*E* and 4*F*, Movie 1, Movie 2). Among three independent MD simulations, the maximum RMSD value of EST in DmNobo[WT] was less than ∼6.60 Å (SI *Appendix*, Fig. S6*A*, Fig. S6*B*). In contrast, with the MD simulation of DmNobo[Asp113Ala], the maximum RMSD value was less than ∼9.54 Å (SI *Appendix*, Fig. S6*A*, Fig. S6*B*). These simulation results also support the possibility that hydrogen bonding between Asp113 and EST is required for stable binding of EST to the H-site.

### Evolutionary conservation of Asp113 in Noppera-bo

Previous reports have demonstrated that the *nobo* family of GSTs is found in Diptera and Lepidoptera (18, 30, 31). Amino acid-sequence analysis revealed that all Nobo proteins from 6 dipteran and 13 lepidopteran species have Asp at the position corresponding to Asp113 of DmNobo (Fig. 3*A*, Fig. 3*B*, Fig. 3*D*). An exception is found in Nobo of the yellow fever mosquito *Aedes aegypti*, as the corresponding amino acid residue of *A. aegypti* Nobo is Glu, which also has a carboxyl group in the sidechain similar to Asp. In contrast, no Asp/Glu residue was found at the corresponding position of the DmGSTD/E/T proteins, other than Nobo (Fig. 3*C*, Fig. 3*E*, Fig 3*F*). Consistent with the amino acid composition, EST did not inhibit the enzymatic activity of the DmGSTE6 or DmGSTE9 recombinant proteins (Fig. 3*G*). These results suggest that Nobo proteins utilize Asp113 to recognize their target compounds as a common feature and that Asp113 serves a biological role.

### Asp113 is essential for *Drosophila melanogaster* embryogenesis

Finally, we examined whether Asp113 is essential for any *in vivo* biological function of DmNobo. We utilized a CRISPR-Cas9-based knock-in strategy to generate a *nobo* allele encoding an Asp113Ala point mutation (*nobo^3×FLAG-HA-D113A^*). We found that no trans-heterozygous mutant *D. melanogaster* with *nobo^3×FLAG-HA-D113A^* and the complete loss-of-*nobo*-function allele (*nobo^KO^*) (18) survived to the adult stage (Table 1). By performing a detailed developmental-stage analysis, we identified no first-instar larvae or later-staged insects with the *nobo^3×FLAG-HA-D113A^*/*nobo^KO^*genotype. These results indicate that the *nobo^3×FLAG-HA-D113A^*/*nobo^KO^*genotype is embryonic lethal. We also found that *nobo^3×FLAG-HA-D113A^*/*nobo^KO^* embryos exhibit an undifferentiated cuticle phenotype (Fig. 5*A*, Fig. 5*B*) and a failure of head involution (Fig. 5*C*, Fig. 5*D*). These phenotypic characteristics were very similar to the feature of Halloween mutants, such as *nobo^KO^*/*nobo^KO^* homozygotes (18). We confirmed that the protein level of Nobo^3×FLAG-HA-D113A^ was comparable to that of Nobo^3×FLAG-HA-WT^ (Fig. 5*E*, Fig. 5*F*), suggesting that the phenotypes were due to loss of protein function, but not impaired gene expression. Taken together, these results show that Asp113 of DmNobo serves a biological function in normal development from the embryonic stage to the adult stage.

**Figure 5.**
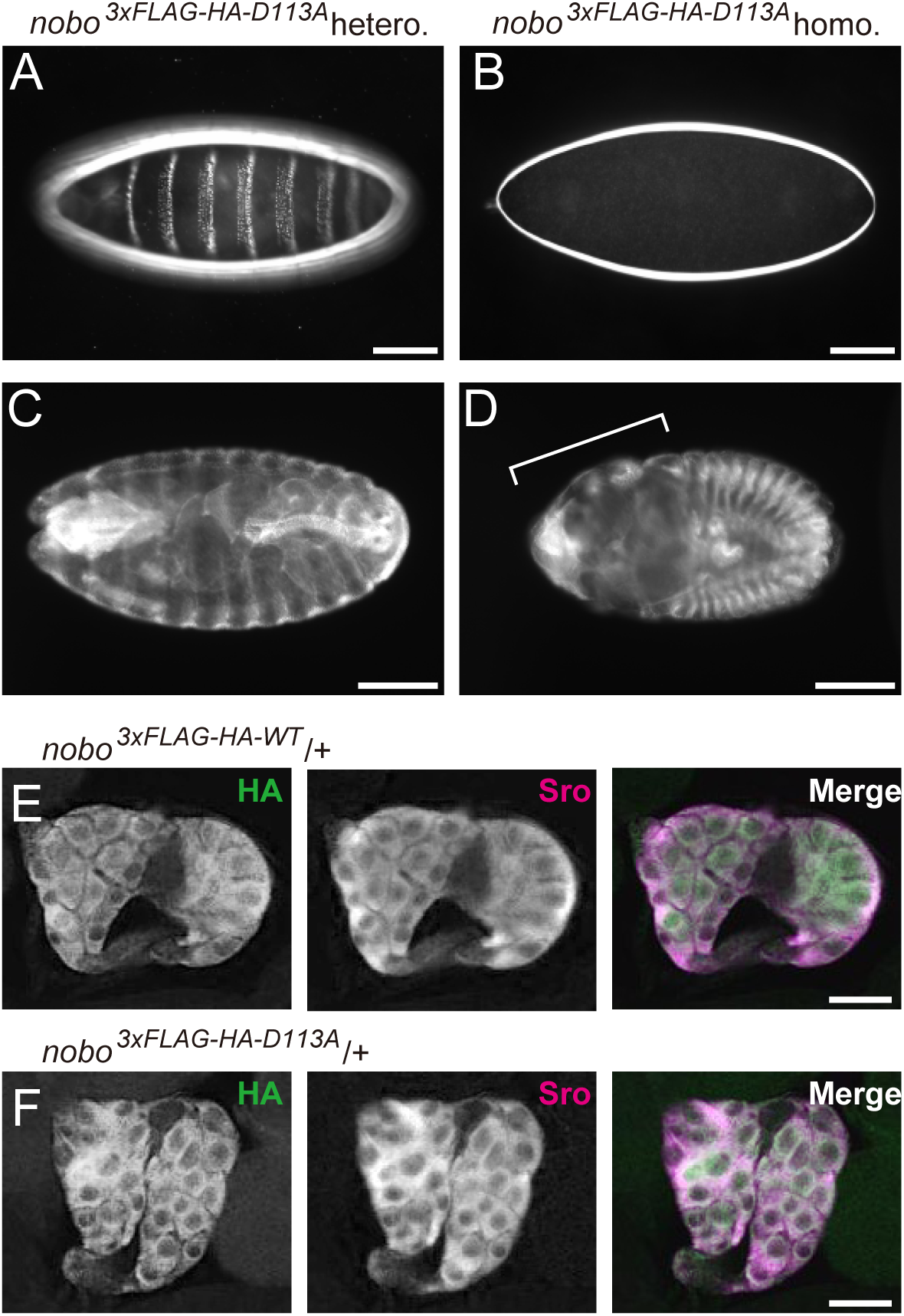
*in vivo* analyses of Asp113Ala. (A, B) Dark-field images of embryonic cuticles from *nobo^3^*^×*FLAG-HA-D113A*^ heterozygotes (*nobo3*×*FLAG-HA-D113A*/*CyO*; A) and homozygotes (*nobo3*×*FLAG-HA-D113A*/*nobo3*×*FLAG-HA-D113A*; B) (C, D) Anti-FasIII antibody staining to visualize overall embryo morphologies. (C) *nobo^3^*^×*FLAG-HA-D113A*^ heterozygotes. (D) *nobo^3^*^×*FLAG-HA-D113A*^ homozygotes. The bracket indicates defective head involution. (E, F) Immunohistochemistry for the ring glands from *nobo^3^*^×*FLAG-HA-D113*^-heterozygous (E) and *nobo^3^*^×*FLAG-HA-D113A*^-heterozygous (F) third-instar larvae. Green and magenta represent the immunostaining observed with anti-HA and anti-Shroud (Sro) antibodies, respectively. Sro was detected as a marker of the prothoracic gland. Scale bars: 100 μm for A–D and 50 μm for E and F

**Table 1.** Viability of *nobo^3×FLAG-HA-D113A^*/*nobo^KO^* knock-in animals

| Background | Knock-in gene | Mating<br>*w*; *nobo*<sup>KO</sup>/<br>CyO-GFP (female) × | Number of adults<br>Cy- (Cy+)<sup>*</sup> | Number of first instar larvae without GFP (with GFP) |
| --- | --- | --- | --- | --- |
| *nobo*<sup>KO</sup> | *nobo*<sup>3×FLAG-HA-WT</sup> | *w*; *nobo*<sup>3×FLAG-HA-WT</sup>/<br>CyO-GFP (male) | 83 (172) | N.D.<sup>†</sup> |
|  | *nobo*<sup>3×FLAG-HA-D113A</sup> | *w*; *nobo*<sup>3×FLAG-HA-D113A</sup>/<br>CyO-GFP (male) | 0 (187) | 0 (157) |

<sup>\*</sup> Cy- and Cy+ indicate animals with straight wings and curly wings, respectively.
<sup>†</sup> N.D. indicates “not determined”.

## Discussion

In this study, we employed an integrated experimental approach, involving *in silico, in vitro*, and *in vivo* analyses to unravel the structure–function relationship of the ecdysteroidogenic GST protein, Nobo. GSTs are widely expressed in all eukaryotes and are also massively duplicated and diversified (24). Among them, the *nobo* family of GST proteins is strictly required for ecdysteroid biosynthesis in insects. Importantly, the lethality of *nobo* mutation in *D. melanogaster* is rescued by overexpressing *nobo* orthologues, but not by overexpressing non-*nobo*-type *gst* genes involved in detoxification and pigment synthesis (18). This fact strongly indicates that, when compared to canonical GSTs, Nobo proteins must possess a unique structural property that make Nobo specialized for ecdysteroid biosynthesis. Regarding this point, this study is significant in that we found that the unique acidic amino acid, Asp/Glu113, is crucial for the *in vivo* function of Nobo. It should be noted that, besides Asp/Glu113, other amino acids constituting the H-sites are also highly conserved among 21 Nobo proteins (Fig. 3*A*, Fig. 3*B*, Fig. 3*D*). These common features imply that the Nobo proteins might share an identical endogenous ligand for the H-site in the ecdysteroidogenic tissues among the species.

An endogenous ligand for Nobo remains a mystery. This study, however, provides some clues for considering candidates for an endogenous ligand. First, it is very likely that the ligand forms a hydrogen bond with the Oδ/Oε atom of Asp/Glu113, given that the *nobo* Asp113Ala point mutation was embryonic lethal and the complete loss-of-function *nobo* phenocopy in mutant *D. melangaster*. Second, considering the complementary shape between the H-site and EST, it seems reasonable to predict that the endogenous ligand(s) is at least similar in shape to steroids. This prediction is also supported by the fact that Nobo acts in ecdysteroidogenic tissues where steroidal molecules are enriched. One steroid that possesses these features is cholesterol. Evidence from our previous study suggests that *nobo* may be involved in cholesterol transport and/or metabolism in ecdysteroidogenic tissues (17–19). Very interestingly, an MD simulation indeed predicted that cholesterol can stably bind to the H-site of DmNobo via a hydrogen bond between the hydroxyl group of cholesterol (C3 position) and Asp113 of DmNobo (SI *Appendix*, Fig. S7). However, paradoxically, it seems that cholesterol contains no site for a chemical reaction with GSH by DmNobo. It is possible that Nobo might serve as a carrier for the ligand, possibly cholesterol, as several classes of GSTs have been shown to exhibit “ligandin” function to carry and transport specific ligands in cells (32). Currently, we have failed in multiple attempts to detect DmNobo-cholesterol complexes via crystallographic analyses, and further experiments are needed for clarify any interaction between Nobo and cholesterol.

The activities of insect ecdysteroids can be disrupted *in vivo* using chemical agonists and antagonists of the ecdysone receptor, some of which are also utilized as insecticides (33). However, chemical compounds that specifically inhibit ecdysteroid biosynthesis are not available. This study provides the first structural information for guiding the development of efficient Nobo inhibitors, which might serve as seed compounds for new insecticides in the future. However, it should be noted that EST and estrogenic chemical compounds are often recognized as endocrine-disrupting chemicals that can dangerously influence the endocrine systems of wild animals (34). Therefore, while EST is a prominent inhibitor of Nobo, a practical compound that can be utilized as an actual insecticide must display no-estrogenic activity. To consider this problem, it is important to note a difference in the EST-recognition patterns between DmNobo and the mammalian estrogen receptor alpha (ERα) protein (35–38). The details of the EST-ERα interaction were investigated using the crystal structures of human ERα in an EST-bound form (35, 39). In ERα, Glu353 interacts with the O3 atom of EST, Phe404 interacts with the A-ring of EST via a CH/π interaction, His524 interacts with the O17 atom of EST, and hydrophobic residues interact with the steroid nucleus. Each of these recognition patterns were found in DmNobo, except for a hydrogen bond with the O17 atom of EST. Asp113 of DmNobo interacts with the O3 atom of EST as in the case of Glu353 of ERα, and DmNobo utilizes a Cys residue of GSH for an SH/π interaction with the A-ring of EST. However, no interaction was found between O17 of EST and residues of the H-site of DmNobo (SI *Appendix*, Fig. S8). Given this difference, we expect that a Nobo-specific, non-estrogenic chemical compound can be developed. Currently, we are pursuing large-scale computational calculations to select chemical compounds that satisfy those conditions and an *in vitro* enzymatic assay to examine DmNobo inhibition.

We emphasize that this report is the first to describe the physical interactions between a Halloween protein and a potent inhibitor at the atomic level. Our interdisciplinary approach will also be applicable for Nobo proteins other than *D. melanogaster*, such as disease vector mosquitos and the agricultural pest moths, and might be a viable strategy for developing new insecticides useful for human societies.

## Materials and Methods

Additional methods are presented in the SI *Appendix*.

### Protein expression, purification, and *in vitro* enzymatic analysis

Recombinant DmNobo was expressed with the pCold-III plasmid vector (TaKaRa Bio) in *Escherichia coli* strain BL21(DE3) (Promega) and purified via glutathione-affinity column chromatography, followed by size-exclusion column chromatography. *In vitro* GST assays and GST activity-inhibition assays using 3,4-DNADCF were performed as described previously (23).

### Crystallization

DmNobo_Apo crystals were obtained using a crystallization buffer containing 42.5% (v/v) polypropylene glycol 400 (PPG400) and 100 mM Bis-Tris (pH 6.4). Crystals of DmNobo-ligand complexes were prepared by the soaking method. DmNobo crystals were soaked for 6 h in an artificial mother liquor (42.5% [w/v] PPG400 and 100 mM Bis-Tris [pH 6.4]) containing 10 mM EST, with or without 1 mM GSH.

### Crystal structure determinations

Crystals were scooped with a cryo-loop (MiTeGen) and flash frozen in liquid nitrogen. Diffraction data were collected at 100 K at beamlines BL-1A and BL-5A in the Photon Factory, Tsukuba, Japan, and at beamline X06SA in the Swiss Light Source, Villigen, Switzerland. The diffraction datasets were automatically processed and scaled using XDS (40), POINTLESS (41), and AIMLESS (42) on PReMo (43). The crystallographic statistics are summarized in SI *Appendix*, Table S1. Phases for the DmNobo_Apo_1 crystal were determined by the molecular replacement method with MOLREP (44) using DmGSTE7 (PDB ID = 4PNG) as an initial model. Molecular replacement calculations for the other structures were performed using DmNobo_Apo_1 as an initial model. Crystallographic refinements were carried out using REFMAC5 and PHENIX.REFINE (45, 46). Models were manually built by COOT (47). The volume of the H-site of DmNobo was calculated using 3V (48). All molecular graphics presented in this manuscript were prepared with the PyMOL Molecular Graphics System, version 1.7.6 (Schrödinger, LLC).

### SPR assay

*K*d values were determined by SPR using a Biacore T200 instrument and a CM5 sensorchip (GE Healthcare) at 25°C. DmNobo[WT] or DmNobo[Asp113Ala] was used as a ligand, and EST was used as an analyte in a buffer containing phosphate buffered saline (PBS) (10 mM KH_2_PO_4_-Na_2_HPO_4_, pH 7.4, 137 mM NaCl, and 2.7 mM KCl) with 1% dimethyl sulfoxide (DMSO) in the presence or absence of 1 mM GSH as a running buffer. The *K*d values of the ligands and analyte were evaluated with Biacore T200 Evaluation Software from triplicate assays.

### FMO calculations

*Ab initio* FMO calculations (49–51) were performed with the DmNobo crystal structures. First, hydrogen atoms were added to the crystal structures, and their energy minimizations were performed. IFIEs were then calculated by the FMO method at the MP2/6-31G* level for the energy-minimized DmNobo models that were fragmented into amino acid and ligand (EST) units. The obtained IFIEs were decomposed into four energy components (the ES, EX, CT+mix, and DI components) using paired interaction-energy decomposition analysis (PIEDA) (28, 29).

### MD simulations

The structures of DmNobo[WT]_EST-GSH and DmNobo[Asp113Ala]_EST-GSH were processed to assign bond orders and hydrogenation. The ionization states of EST and GSH suitable for pH 7.0 ± 2.0 were predicted using Epik (52), and H-bond optimization was conducted using PROPKA (53). Energy minimization was performed in Maestro using the OPLS3 force field (54). Preparation for MD simulation was conducted using the Molecular Dynamics System Setup Module of Maestro (Schrödinger). DmNobo[WT]_EST-GSH and DmNobo[Asp113Ala]_EST-GSH, which were subjected to energy minimization, were placed in an orthorhombic box with a buffer distance of 10 Å to create a hydration model, using the TIP3P water model (55). NaCl (0.15 M) was added as the counterion to neutralize the system. The MD simulations were performed using Desmond ver. 2.3 (Schrödinger) (56). The cut-off radii for the van der Waals and electrostatic interactions, and the time step, initial temperature, and pressure of the system were set to 9 Å, 2.0 fs, 300 K, and 1.01325 bar, respectively. The sampling interval during the simulation was set to 10 ps. Finally, we performed MD simulations with the NPT ensemble for 100 ns.

### Phylogenetic analysis

Twenty one amino acid sequences of DmNobo or *Bombyx mori* Nobo orthologues were obtained from NCBI non-redundant protein database (57) and the Uniprot Knowledgebase (58).

For the phylogenetic analysis of insect GSTD/E/T proteins, previously described amino acid sequences were obtained from the Uniprot Knowledgebase, NCBI protein database, MonarchBase (18, 58–60). Multiple alignment of 503 amino acid sequences was performed with COBALT (61), and the sequence alignment was used for cluster-analysis with CLANS (62). A major cluster included 372 amino acid sequences of GSTD/E/T and other GST proteins (*SI Appendix*, Table S5). A phylogenetic tree was drawn with COBALT using the 372 GSTs with a neighbor-joining algorithm. We identified 371 sequences with a Grishin-sequence difference of 0.9, which included 151 GSTDs, 178 GSTEs, and 42 GSTTs. The GSTEs included 21 Nobo proteins. To calculate the amino acid frequencies, the obtained alignment was manually edited based on the known crystal structures, using Jalview (63). The amino acid frequencies were calculated and illustrated using WebLOGO version 3.7.4 (64), and colored using the “Chemistry (AA)” scheme.

### Transgenic *D. melanogaster* insects and genetics

The generation of *D. melanogaster* knock-in flies was performed as described in the *SI materials*. We found that *nobo^3×FLAG-HA-WT^*^−^homozygous flies were fully viable, whereas *nobo^3×FLAG-HA-D113A^-*homozygous flies displayed embryonic lethality. We utilized *nobo^3×FLAG-HA-D113A^*-heterozygous and -homozygous embryos for cuticle preparation and immunostaining. To formally rule out the possibility that the embryonic lethality was due to anonymous deleterious mutations other than *nobo^3×FLAG-HA-D113A^*, we counted the number of trans-heterozygous flies with a *nobo* knock-out (*nobo^KO^*) from a previous report (18), as follows. Heterozygous *nobo^3×FLAG-HA-WT^*, *nobo^3×FLAG-HA-D113A^*, and *nobo-*knock-out (*nobo^KO^*) alleles were balanced with *CyO* carrying *Actin5C:gfp* cassette (*CyO-GFP)*. Either *nobo^3×FLAG-HA-WT^/CyO-GFP* flies or *nobo^3×FLAG-A-D113A^/CyO-GFP* flies were crossed with *nobo^KO^/CyO-GFP* flies. The trans-heterozygous flies (*nobo^3×FLAG-HA-WT^/nobo^KO^*or *nobo^3×FLAG-HA-D113A^/nobo^KO^*) should exhibit no GFP signals. We found that GFP-negative *nobo^3×FLAG-HA-WT^/nobo^KO^*embryos hatched normally and developed into adults without any abnormalities, whereas *nobo^3×FLAG-HA-D113A^/nobo^KO^*embryos did not.

### Cuticle preparation and immunostaining

Embryonic cuticle preparation was performed as previously described (65). Immunostaining for whole-mount embryos was conducted as previously described (18). A mouse anti-FasIII monoclonal antibody 7G10 (Developmental Studies Hybridoma Bank, University of Iowa, USA; 1:20 dilution) and an anti-mouse IgG antibody conjugated with Alexa488 (Life Technologies; 1:200 dilution) were used for immunostaining the embryos. For immunostaining of the brain-ring gland complex in third-instar larvae, we first crossed *nobo^3×FLAG-HA-WT^* homozygous females or *nobo^3×FLAG-HA-D113A^*/*CyO-GFP* females with Oregon-R wild type males. Third-instar larvae of the heterozygous offspring (*nobo^3×FLAG-HA-WT^/+* or *nobo^3×FLAG-HA-D113A^/+*) were dissected and then immunostained as previously described (66). The antibodies used for the brain-ring gland complex included a rat anti-HA high-affinity monoclonal antibody (3F10, 1:20 dilution; Roche), a guinea pig anti-Shroud antibody (67) (1:200 dilution), an anti-rat IgG antibody conjugated with Alexa488 (1:200 dilution; Life Technologies), and an anti-guinea pig IgG antibody conjugated with Alexa555 (1:200 dilution; Life Technologies). Fluorescence images were obtained using an LSM700 microscope (Carl Zeiss).

## Supporting information

Supplementary text Figs. S1 to S10, Tables S1 to S5, Legends of Movie 1 and 2, References for SI reference citations

PDB Validation Report ID 6KEL

PDB Validation Report ID 6KEM

PDB Validation Report ID 6KEN

PDB Validation Report ID 6KEO

PDB Validation Report ID 6KEP

PDB Validation Report ID 6KEQ

PDB Validation Report ID 6KER

## Acknowledgements

We thank Teruki Honma and Chiduru Watanabe at RIKEN for discussions regarding the FMO calculations; Tamie Katsuta, Hiroyuki Matsumaru, and Ryuichi Kato for setting up crystal-screening conditions; Dr. Yusuke Yamada for managing the automated data collection at the Photon Factory; Ayaka Harada, Akira Shinoda, and Miki Senda for data collection at the Swiss Light Source; and the Developmental Studies Hybridoma Bank (University of Iowa) for providing us with antibodies. We are also grateful to Yoshiaki Nakagawa and Hajime Ono for their critical reading of the manuscript; and Tetsuo Nagano for his continuous encouragement and kind advice. We acknowledge the Paul Scherrer Institut (Villigen, Switzerland) for providing synchrotron radiation beamtime at beamline X06SA of the Swiss Light Source (proposal numbers 20181219 and 20181299). This work was supported in part by a KEK Postgraduate Research Student fellowship to K.I. This work was also supported by KAKENHI (grant numbers 15K14719 and 18K19163) to R.N. and by the Private University Research Branding Project. In addition, this work was supported by the Platform Project for Supporting in Drug Discovery and Life Science Research (Platform for Drug Discovery, Informatics, and Structural Life Science) and the Basis for Supporting Innovative Drug Discovery and Life Science Research from Japan Agency for Medical Research and Development (AMED) under Grant Number 18am0101113j0002. This work was performed under the approval of the Photon Factory Program Advisory Committee (proposal number 2018G025).

## Footnotes

Author contributions: K.Ko., K.I., Y.F., F.Y., T.S., and R.N. designed the research; K.Ko., K.I., K.M., F.Y., and T.S. performed the X-ray crystallographic analysis; K.I., K.M., S.E., R.A., H.K., T.O. Y.F., and H.I. performed the enzymatic assays and analyzed the data; K.Ko., K.I., and R.A. performed the surface plasmon-resonance assays; R.Y. and T.H. performed MD simulations; K.Ka. and K.F. performed FMO calculations; R.A., Y.S.N., A.N. and R.N. performed experiments with the fruit flies; and K.Ko., K.I., R.Y., K.F., F.Y., T.S., and R.N. wrote the manuscript.

The authors declare no conflict of interest.

This article is a PNAS Direct Submission.

Data deposition: The X-ray data and coordinates presented in this paper were deposited in the Protein Data Bank (https://pdbj.org/) under the following PDB IDs: 6KEL, 6KEM, 6KEN, 6KEO, 6KEP, 6KEQ, and 6KER.

