## Supplementary text Figs. S1 to S10, Tables S1 to S5, Legends of Movie 1 and 2, References for SI reference citations for "An integrated approach unravels a crucial structural property for the function of the insect steroidogenic Halloween protein Noppera-bo"

**This PDF file includes:**

Supplementary text

Figs. S1 to S10

Tables S1 to S5

Legends of Movie 1 and 2

References for SI reference citations

SI APPENDIX MATERIALS AND METHODS

**Protein expression and purification**

The protein-expression plasmid pCold-III (Takara Bio Inc., Kusatsu, Japan) was used to express recombinant GST proteins in *E. coli*. Coding sequences (CDSs) of *Drosophila melanogaster nobo* (*CG4688, Dmnobo*), *gste6* (*CG17530, Dmgste6*), and *gste9* (*CG17534, Dmgste9*) were amplified by the polymerase chain reaction (PCR) using complementary DNA derived from *D. melanogaster* larvae. The primers used for PCR were nobo-Fwd (5′-CAGTCATATGATGTCTCAGCCCAAGCCGATTTTG-3′), nobo-Rev (5′-CTCGAGCTACTCCACCTTCTCGGTGACTACCG-3′), GSTe6-Fwd (5′-CATATGATGGTGAAATTGACTTTATACGG-3′), GSTe6-Rev (5′-TCTAGATCATGCTTCGAATGTGAAATT-3′), GSTe9-Fwd (5′-CATATGATGGGAAAATTAGTACTGTACGG-3′), and GSTe9-Rev (5′-TCTAGATTACACAATCTTTGTGATCTTCG-3′). The *nobo* CDS was subcloned between the *Nde* I and *Xho* I restriction enzyme sites in pCold-III to generate the pCold-III_DmNobo[WT] vector. The *gste6* and *gste9* CDSs were subcloned between the *Nde* I and *Xba* I sites in pCold-III. It should be noted that pCold-III added a translation enhancing element (MNHKV) at the N-terminus of each of DmNobo, DmGSTE6 and DmGSTE9 proteins.

An expression vector for DmNobo[Asp113Ala] was constructed by inverse-PCR-based site-directed mutagenesis. The entire pCold-III_DmNobo[WT] plasmid was amplified by inverse PCR using a KOD-Plus-Mutagenesis Kit (Toyobo Co., Ltd, Osaka, Japan) using the oligonucleotides, 5′-CCAGTGATTTTATGTCGGCGATTGTCCGCC-3′ and 5′-CACGTCGGAACAAAAAGGAGCATTCGAAGA-3′, as amplification primers. The *E. coli* strain DH5α was transformed with the *Dpn* I-digested PCR product. The plasmids were purified using a FastGene Plasmid Mini Kit (NIPPON Genetics Co., Ltd., Tokyo, Japan). Those DNA sequences were confirmed by Sanger sequencing with one of the following sequencing primers: 5′-ACGCCATATCGCCGAAAGG-3′ or 5′-GGCAGGGATCTTAGATTCTG-3′.

DmNobo, *D. melanogaster* GSTE6 (DmGSTE6) and *D. melanogaster* GSTE9 (DmGSTE9) were expressed in the *E. coli* strain BL21(DE3) (Merck, Darmstadt, Germany) and purified via GSH-affinity column chromatography, followed by size-exclusion column chromatography. *E. coli* BL21(DE3) cells were transformed with pCold-III_DmNobo, and then the transformed cells were cultured in LB medium supplemented with 50 µg/mL ampicillin at 37°C. When the OD_600_ of the culture reached approximately 0.6, protein expression was induced with 0.3 mM isopropyl β-D-1-thiogalactopyranoside. The *E. coli* cells were cultured at 18°C overnight and then harvested. The harvested cells were suspended in lysis buffer (300 mM NaCl, 25 mM Tris-HCl, 1 mM CHAPS, 1 mM DTT, pH 8.0) and lysed for 2 min by sonication using a VP-305 Ultra 5 Homogenizer (TAITEC), using an output of 7 and a duty of 40%. The lysate was fractionated by centrifugation at 15,000 × *g* for 30 min at 4°C, and the supernatant was applied to a GSH-affinity column containing a 10-ml bed volume of glutathione Sepharose 4B (GE Healthcare, Little Chalfont, United Kingdom). After the column was washed with lysis buffer, the proteins were eluted with 50 ml of elution buffer (140 mM NaCl, 25 mM Tris-HCl [pH 8.0], 1 mM CHAPS, 1 mM DTT, 10 mM GSH). The eluent was concentrated to 2 ml and fractionated with a Superdex200 10/300 size-exclusion column (GE Healthcare) connected to an ÄKTA FPLC system (GE Healthcare). The column was equilibrated with a buffer (150 mM NaCl, 25 mM Tris-HCl [pH 8.0], 1 mM DTT), and proteins were eluded with the same buffer at a flow rate of 0.2 mL/min. The peak fractions were concentrated to 15 mg/ml and stored at -80°C. The protein concentrations of DmNobo, DmGSTE6, and DmGSTE9 was measured with a NanoDrop ND-1000 spectrophotometer (Thermo Fisher Scientific, Massachusetts, United States of America) using extinction coefficients (ε_280_) of 0.671•M^–1^•cm–1, 1.274•M^–1^•cm^–1^, and 1.128•M^–1^•cm^–1^, respectively.

***In vitro* GST assay**

*In vitro* GST assays with 3,4-DNADCF were performed as described previously (1). The stock solutions of DmNobo[WT] and DmNobo[Asp113Ala] were 200 ng/mL each in solution A (2 mM GSH, 100 mM sodium phosphate buffer [pH 6.5], 0.01% Tween 20). Decreasing concentrations of DmNobo[WT] and DmNobo[Asp113Ala], ranging from 200 ng/mL to 0.19 ng/mL, were prepared by 2-fold serial dilution with solution A. The DmNobo dilution series was mixed with an equal volume of solution B (100 mM sodium phosphate buffer [pH 6.5] with 2 µM 3,4-DNADCF in 0.2% DMSO as a co-solvent) in each well of a 96-well plate to initiate the catalytic reaction of DmNobo. The GSH-conjugated product was excited at 485 nm, and the fluorescence intensity at 535 nm (*F*_measured_) was measured every 30 s for 20 min with an infinite 200 PRO instrument (Tecan, Zurich, Switzerland). The fluorescence intensity (*F_t_*) in the reaction mixture without DmNobo (*F*_bg_) was subtracted as the background signal (*F*_t_ = *F*_measured_ – *F*_bg_). The maximum fluorescence intensity (*F*_max_) was the fluorescence intensity that was reached as a plateau. The amount of product in each well (*P*_t_) at the measured time (*t*) was calculated as *P*_t_ (µmol) = *F*_t_/*F*_max_ × 200 µL × 1 µmol/L. The rate of product formation (*P*_rate_, µmol/min) was obtained by linear-least-squares fitting between *P*_t_ and *t*. The specific activity of DmNobo (µmol/min/mg-protein) was defined as *P*_rate_/[protein concentration]. The assay was performed in triplicate.

**GST activity-inhibition assay**

EST was dissolved in DMSO to a concentration of 2.5 mM. The 2.5 mM EST solution was diluted to 50 µM EST in solution C (2 mM GSH, 100 mM sodium phosphate buffer, 0.01% Tween 20, 2% DMSO, and 25 ng/mL DmNobo[WT], 25 ng/mL DmNobo[Asp113Ala], 17.5 ng/mL DmGSTE6, or 150 ng/mL DmGSTE9 [pH 6.5]). A dilution series of EST, ranging from 50 µM to 0.19 µM, was prepared by 2-fold serial dilution with solution C. One hundred microliters of each EST solution in the dilution series was mixed with an equivalent amount of solution B (100 mM sodium phosphate buffer [pH 6.5] and 2 µM 3,4-DNADCF in 0.2% DMSO) in each well of a 96-well plate. *F*_measured_ values were measured after 3 min, as described in the “*In vitro* GST assay” section. The fluorescence intensity detected in the absence of EST and DmNobo (*F*_bg_) was subtracted as the background in all experiments (*F* = *F*_measured_ – *F*_bg_). *F* at 0 s (*F*_0_) was subtracted from *F* at the measured time (s) (*F*_t_ = *F* – *F*_0_).

The relative activity was calculated as *F*_30_[I]_/*F*_30_[0]_, where [I] and [0] indicate the EST concentrations. The relative activity was plotted against each EST concentration. A fitting curve was calculated based on a plot generated from the following equation when IC_50_ and Hill constant (n) were approximated as 1.00 and 1.00, respectively, using KaleidaGraph version 4.5.1 (Synergy Software, Reading, United States of America):

Relative activity (%) = 1/(1 + {[EST]/IC_50_)^n^}) × 100

The IC_50_ value was estimated based on the fitting curve.

**Phylogenetic analysis**

Nineteen amino acid sequences of DmNobo or *Bombyx mori* Nobo orthologues were found using BLASTP (2) to search the NCBI non-redundant protein database. In addition, a Nobo orthologue in *H. armigera* was found in the Uniprot Knowledgebase. The accession numbers were XP_021192638.1 for *Helicoverpa armigera* GSTE14-like isoform X2, A0A2W1BRE1 for *H. armigera* uncharacterized protein, XP_022126447.1 for *Pieris rapae* GSTE14-like, XP_022837694.1 for *Spodoptera litura* GSTE14-like isoform X2, PCG75296.1 for *Heliothis virescens* hypothetical protein B5V51_11931, XP_013196516.1 for *Amyelois transitella* GST1, XP_001658748.2 for *Aedes aegypti* GSTE14, XP_319963.1 for *Anopheles gambiae* GSTE8, KXJ68754.1 for *Aedes albopictus* hypothetical protein RP20_CCG001852, ETN60212.1 for *Anopheles darlingi* GSTE, KFB39334.1 for *Anopheles sinensis* AGAP009190-PA-like protein, XP_001868776.1 for *Culex quinquefasciatus*, KOB78695.1 for *Operophtera brumata* GST, AIL29314.1 for *Cnaphalocrocis medinalis* GSTE5 partial region, XP_014368559.1 for *Papilio machaon* GSTE14-like, XP_013137131.1 for *Papilio polytes* GST1-1-like, NP_001299034.1 for *Papilio xuthus* GST1-1, NP_001292431.1 for an uncharacterized protein *Plutella xylostella*, ABY66602.1 for *B. mori* GSTE14, and OWR47941.1 for *Danaus plexippus*. Two *nobo* orthologues were found for *H. armigera* in the database.

For phylogenetic analysis of insect GSTD/E/T proteins, previously described amino acid sequences were obtained from the Uniprot Knowledgebase, NCBI protein database, and MonarchBase (3–6). Amino acid sequences (503) were aligned with COBALT (7), and the resulting sequence alignment was used for cluster analysis with CLANS (8). A major cluster included 372 amino acid sequences, including those of GSTD/E/T proteins and other GST proteins (SI *Appendix*, Table S5). A phylogenetic tree was drawn with COBALT, using the 372 GSTs and a neighbor-joining algorithm. We identified 371 sequences with a Grishin-sequence difference of 0.9, including 151 GSTDs, 178 GSTEs, and 42 GSTTs. We also identified 21 Nobo proteins among the GSTEs.

To calculate the amino acid frequencies, the obtained alignment was manually edited based on the known crystal structures, using Jalview (9). The amino acid frequencies were calculated and illustrated with WebLOGO version 3.7.4 (10).

**Surface plasmon-resonance assay**

Surface plasmon resonance was measured at 25°C using Biacore T200 instrument with a CM5 sensor chip (GE Healthcare). DmNobo[WT] or the DmNobo[Asp113Ala] protein was used as a ligand, and EST was used as an analyte in PBS containing 1% DMSO, in the presence or absence of 1 mM GSH as a running buffer.

The Biacore T200 system with a CM5 sensor chip was filled with the running buffer. The ligands were immobilized on the activated CM5 sensor chip in an acetate buffer (pH 5.0) using a purchased amine-coupling kit (GE Healthcare) to reach 6,500 resonance units. The same process was performed in the absence of proteins in one lane on the chip as a background lane.

An EST dilution series was prepared by serial dilution. An EST stock solution (100 mM EST in DMSO) was diluted with running buffer to a concentration of 20 µM. The 20 µM EST solution was serially diluted by two thirds with running buffer seventeen times, and the running buffer in the absence of EST was used as the 0-µM EST sample. The analyte was flowed onto the sensor chip for 60 s and allowed to dissociate for 180 s.

EST concentrations of 20.000, 13.333, 8.889, 5.926, 3.951, 2.634, 1.756, 1.171, 0.780, 0.520, 0.347, 0.231, 0.154, 0.103, 0.069, 0.046, 0.030, 0.020, and 0.014 µM were used to calculate its *K*d. The background was subtracted from the sensorgrams of the proteins-immobilized lanes (sensorgrams shown in Fig. 3*B*). The *K*d values of EST for DmNobo[WT] and DmNobo[Asp113Ala] were evaluated with Biacore T200 Evaluation Software, using data from triplicate assays.

**Crystallization**

The Protein Crystallization System (11) was used for the initial crystallization screening of DmNobo (16). In total, 384 conditions were examined using the Crystal Screen 1 & 2, Index, PEG/Ion, or PEG/Ion 2 kits from Hampton Research (CA, USA), or the Wizard I & II kit from Molecular Dimensions (Suffolk, United Kingdom). DmNobo was crystallized at 20°C in the presence of 25% (w/v) PEG 3350 in 100 mM Bis-Tris (pH 5.5; index #42), or 45% (v/v) PPG 400 in 100 mM Bis-Tris (pH 6.5; index #58). The crystallization conditions were optimized by changing the pH and the concentration of the precipitation agent, resulting in two types of crystals, DmNobo I and II. DmNobo I crystals were obtained from a buffered solution containing 27.5% (w/v) PEG 3350 in 100 mM MES-NaOH (pH 5.4), and DmNobo II crystals were obtained from a buffered solution containing 42.5% (v/v) PPG400 in 100 mM Bis-Tris (pH 6.4). Crystals of substrate complexes were prepared by soaking the DmNobo II crystals in an artificial mother liquor (42.5% [w/v] PPG 400 in 100 mM Bis-Tris [pH 6.4]) containing 10 mM EST, with or without 1 mM GSH, for 6 h.

**Crystal structure determination and analysis**

Crystals were picked up with proper size of MicroLoops (MiTeGen, New York, USA), flash frozen in liquid nitrogen, and packed in Uni-pucks (Molecular Dimensions). Diffraction data were collected at beamline BL-1A in the Photon Factory (Tsukuba, Japan) for DmNobo_Apo_1 crystals and at beamline X06SA in the Swiss Light Source for DmNobo_Apo_2 and other complexes. The diffraction datasets collected at the Photon Factory were automatically processed and scaled using XDS (12), POINTLESS (13), and AIMLESS (14) on PReMo (15), and those collected at the Swiss Light Source were processed and scaled using XDS and AIMLESS. Crystallographic statistics are summarized in SI *Appendix* Table S1.

Phases for DmNobo_Apo_1 data were determined by the molecular replacement (MR) method with MOLREP (16) using the crystal structure of DmGSTE7 (PDB ID = 4PNG) as a search model. Other crystal structures were determined by the MR method using the crystal structure of DmNobo_Apo_1 as a search model. Molecular models were initially refined with REFMAC5 (17). The models were manually built using COOT (18) and further refined with PHENIX.REFINE (19) repeatedly. The C-terminal four residues could not be modeled due to poor electron density. In this study, the crystal structure of DmNobo_Apo_2 was used as the DmNobo_Apo structure when making comparisons with other crystal structures. m*F*o-D*F*c omit-maps for ligands were calculated using PHENIX.REFINE with a simulated annealing protocol. Interactions between DmNobo and GSH or EST was analyzed using PISA (20). The volume of the cavity in DmNobo was calculated using the Channel Finder program in 3V (21), with 4-Å radius for the outer probe and a 1-Å radius for the inner probe. The volumes of GSH or EST were calculated using the Volume Assessor program in 3V, with a 2-Å radius for each probe. The RMSD from a least-squares fitting among the DmNobo structures was calculated with GESAMT (22). Atom pairs within a 4.0-Å distance were defined as making direct contacts. All molecular graphics were prepared using the PyMOL Molecular Graphics System, version 1.7.6 (Schrödinger, NY, USA).

**Construction of *D. melanogaster* knock-in flies**

*D. melanogaster* flies were reared on standard agar-cornmeal medium at 25°C under a 12 h/12 h light/dark cycle. The strain harboring the Asp113Ala point mutation (*nobo^3^**^×FLAG-HA-D113A^*), as well as the control wild-type strain (*nobo^3×FLAG-HA-WT^*) were generated using a CRISPR-Cas9-mediated knock-in strategy (23). Briefly, in each case, the genome of the starter *yw* strain was cut at 2 sites around the *nobo* locus, and then homologous recombination occurred with appropriate plasmids carrying 5′- and 3′-homology arms and an N-terminal 3× FLAG-HA epitope tag. The pDCC6 plasmid was used for simultaneous expression of both the Cas9 gene and guide RNA (gRNA) (24). The following primer pairs were annealed and then ligated to *Bbs* I-digested pDCC6, which led to the production of 3 different gRNA plasmids: 5′-CTTCGTTGGGCTGAGACATTAAGTT-3′ and 5′-AAACAACTTAATGTCTCAGCCCAAC-3′ for Cutter#1; 5′-CTTCGTTACGACGAGCGCAGTCCGC-3′ and 5′-AAACGCGGACTGCGCTCGTCGTAAC-3′ for Cutter#2; and 5′-CTTCGCCGACGTGACAGTGATTTTA-3′ and 5′-AAACTAAAATCACTGTCACGTCGGC-3′ for Cutter#3. The pUC19-based plasmids carrying the homology arms and epitope tags, designated pDonor[KI]-{CG4688_LA}:{3×FLAG/HA/nobo}:{CG4688_RA} and pDonor[KI]-{CG4688_LA}:{3×FLAG/HA/nobo*D113A}:{CG4688_RA}, respectively, were artificially synthesized by VectorBuilder, Inc (Chicago, IL, USA). The entire DNA sequence of each plasmid is shown in SI *Appendix*, Fig. S10 and Fig. S11. To generate the *nobo^3×FLAG-HA-WT^* strain, the Cutter#1, Cutter#2, and pDonor[KI]-{CG4688_LA}:{3×FLAG/HA/nobo}:{CG4688_RA} plasmids were injected into *yw* embryos. To generate the *nobo^3×FLAG-HA-D113A^* strain, the Cutter#1, Cutter#3, and pDonor[KI]-{CG4688_LA}:{3×FLAG/HA/nobo*D113A}:{CG4688_RA} plasmids were injected to *yw* embryos. The proper knock-in strains were identified and characterized, essentially as described previously (25). DNA sequences surrounding the knock-in regions were confirmed by Sanger sequencing.

**FMO calculations**

*Ab initio* FMO calculations (26–28) were performed on the crystal structures of the DmNobo_Apo, DmNobo_EST-GSH, DmNobo_GSH, and DmNobo_EST complexes. While DmNobo is a homodimer, only the monomeric structure was utilized for the FMO calculations. Inter-subunit interactions were therefore neglected in this study. The crystal structures were modified before performing the FMO calculations. First, all crystal water molecules, except for one that interacts with the carbonyl oxygens of Glu in GSH and Pro58, and the Oγ atom of Ser56 of DmNobo (Fig. 2*A*, water in yellow), were deleted from the crystal structures. Second, assignment of the protonation state and the addition of hydrogen atoms were performed using the Protonate 3D function of the Molecular Operating Environment program package (Chemical Computing Group, Montreal, Canada). Then, energy minimization of hydrogen atoms was performed with the Amber10:EHT force field. The protonated states of His55 and His71 were assumed to be positively charged to form hydrogen bonds with GSH. Then, FMO calculations for the monomeric DmNobo structures were performed using ABINIT-MP software (29, 30). The second-order Møller-Plesset perturbation (MP2) (31, 32) method was used with the 6-31G* basis function as a theoretical calculation level; namely, the FMO-MP2/6-31G* level of theory was used. For the FMO calculations, DmNobo proteins and GSH were fragmented into amino acid units at bonds between the C and Cα atoms of the main-chain. Each EST and water molecule was treated as a single fragment. The fragmentation treatment makes it possible to easily calculate the electronic structure of the whole complex and the IFIEs. The obtained IFIEs were further decomposed into four energy components, i.e., the ES, EX, CT+mix, and DI components, using PIEDA (33, 34).

**MD simulations**

The structures of DmNobo[WT]_EST-GSH, DmNobo[Asp113Ala]_EST-GSH, and DmNobo_cholesterol-GSH, were processed to assign bond orders and hydrogenation. The ionization states of EST, cholesterol, and GSH at pH 7.0 ± 2.0 were predicted using Epik (35), and H-bond optimization was conducted using PROPKA (36). Energy minimization was performed in Maestro using the OPLS3 force field (37).

Preparation for MD simulations was conducted using the Molecular Dynamics System Setup Module of Maestro (Schrödinger, NY, USA). DmNobo[WT]_EST-GSH and DmNobo[Asp113Ala]_EST-GSH were subjected to energy minimization and placed in an orthorhombic box with a buffer distance of 10 Å to create a hydration model, and the TIP3P water model (38) was used for the hydration model. NaCl (0.15 M) served as the counterion to neutralize the system.

The MD simulations were performed using Desmond software, version 2.3 (Schrödinger, NY, USA). The cut-off radii for van der Waals and electrostatic interactions, and the time step, initial temperature, and pressure of the system were set to 9 Å, 2.0 fs, 300 K, and 1.01325 bar, respectively. The sampling interval during the simulation was set to 10 ps. Finally, we performed MD simulations using the NPT ensemble for 100 ns.

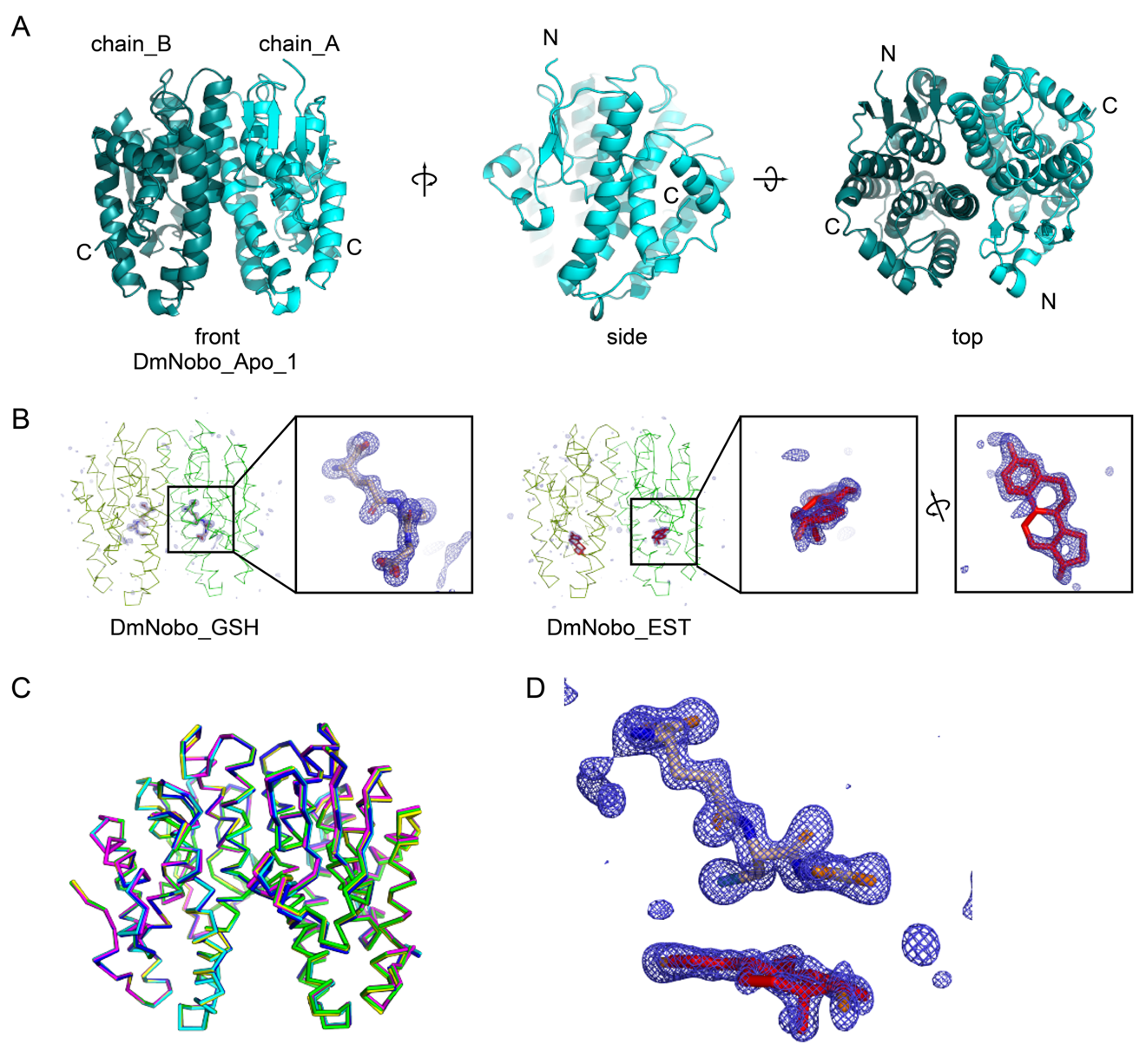

Fig. S1. Overall structures of DmNobo.

(A) The overall structure of DmNobo_Apo_1 is represented as a ribbon diagram. Chains A and B are colored in cyan and turquoise, respectively.

(B) Simulated annealing-omit maps of GSH and EST in DmNobo_GSH and DmNobo_EST, respectively. m*F*o-D*F*c maps (blue) within a 5-Å distance from all ligand atoms are contoured at the 4.0σ level and shown with a blue mesh. A ribbon model of the DmNobo protein (green) and stick models of GSH and EST molecules are shown. Carbon atoms in GSH and EST are colored wheat and red, respectively. Oxygen atoms and nitrogen atoms are colored in green and blue, respectively.

(C) Superimposed crystal structures of DmNobo_Apo_1 (cyan), DmNobo_Apo_2 (blue), DmNobo_GSH (yellow), DmNobo_EST (magenta), and DmNobo_EST-GSH (green). Least square (LSQ) fittings were performed with the Cα atoms of chain A. RMSD values from the LSQ fittings are summarized in Table 1.

(D) Simulated annealing-omit maps of GSH and EST in the DmNobo_EST-GSH structure. m*F*o-D*F*c maps (blue) are contoured at 4.0σ and shown with a blue mesh. GSH and EST molecules are shown as stick models.

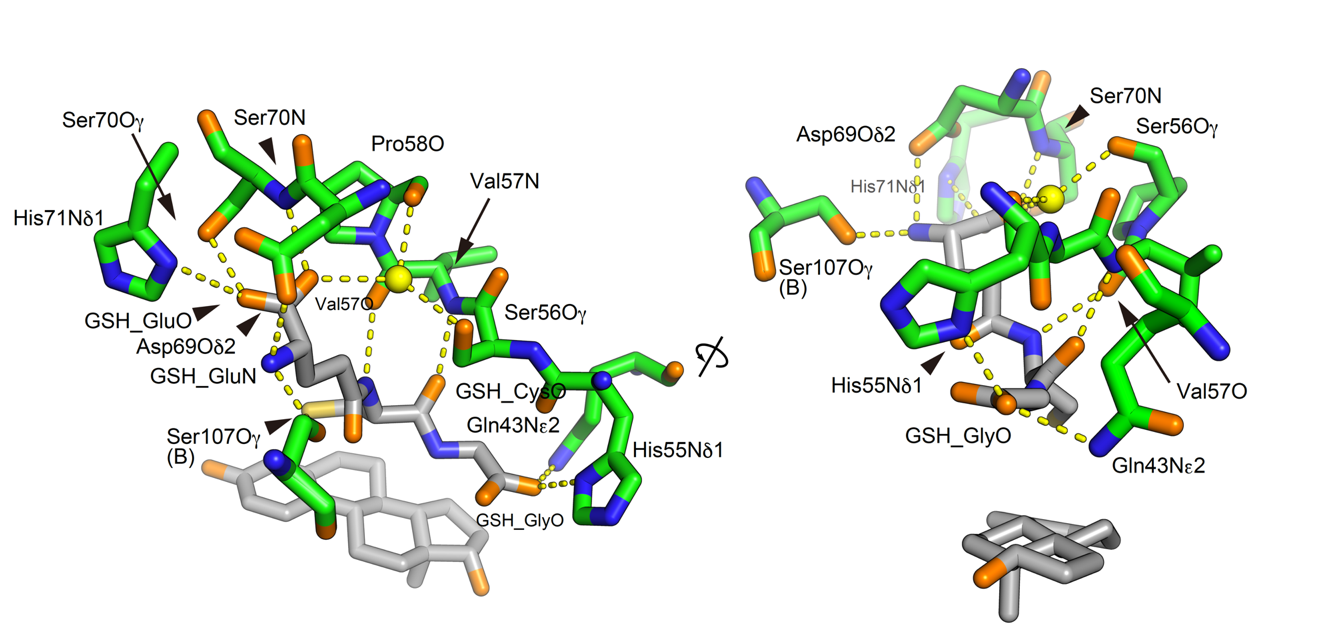

Fig. S2. A hydrogen-bond network between DmNobo and GSH.

Hydrogen bonds are indicated with yellow dashed lines. Carbon atoms of the protein and ligands (GSH, EST) are shown in green and gray, respectively. One of the two conformations of Ser107, in which the Oγ atom is directed towards GSH, is shown.

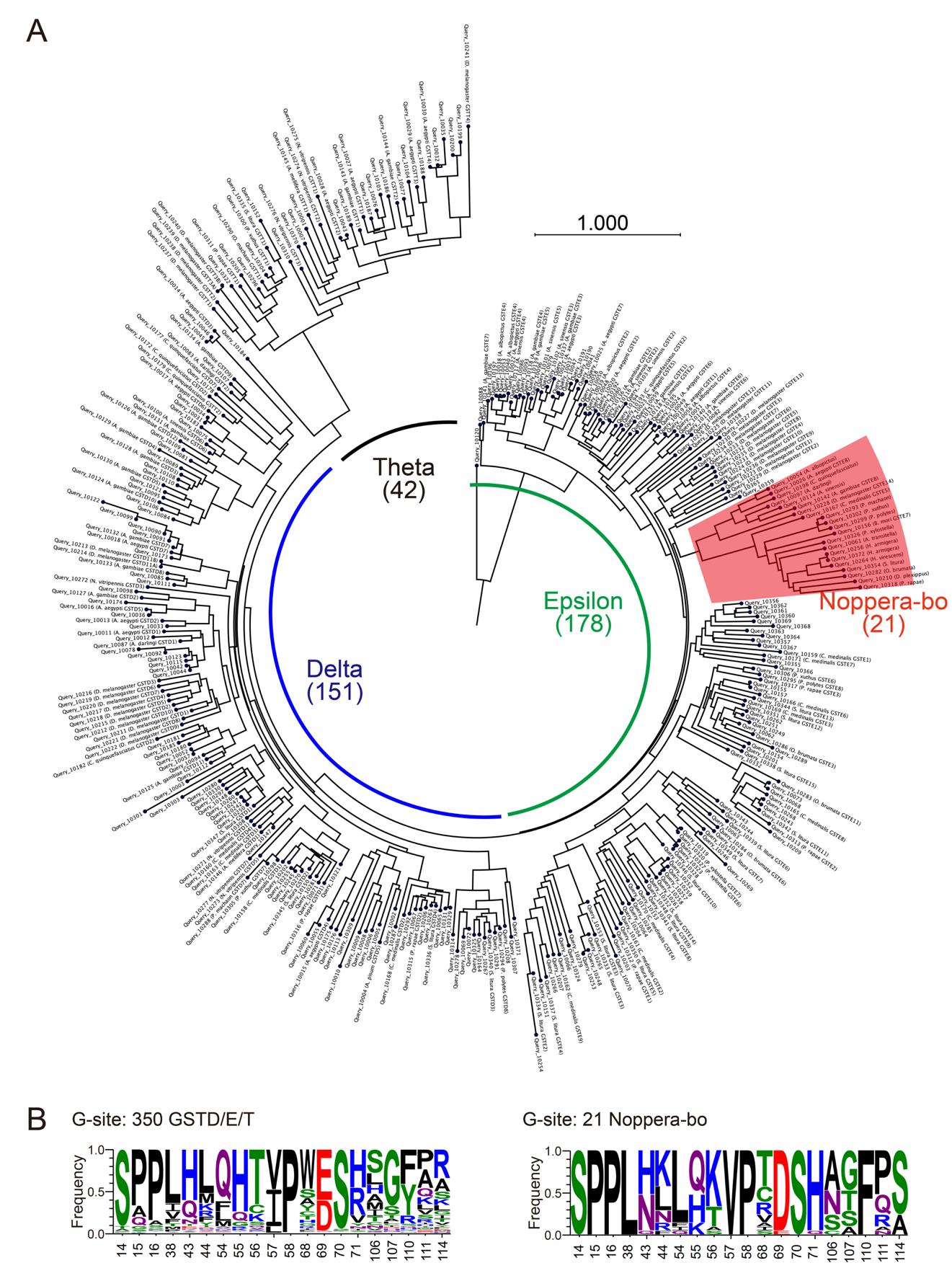

**Fig. S3. Phylogenetic analysis of the delta, epsilon, and theta classes of GST.**

(A) Phylogenetic tree of the insect GSTD/E/T proteins listed in SI *Appendix*, Table S5. A clade of the *nobo* family is highlighted in red.

(B) Frequencies of amino acid residues composing the G-site of 350 GSTD/E/T (left) and 21 Nobo (right) proteins displayed in A. The frequencies were calculated using LOGO (10), and Nobo proteins were excluded from the GSTD/E/T proteins for the frequency calculation.

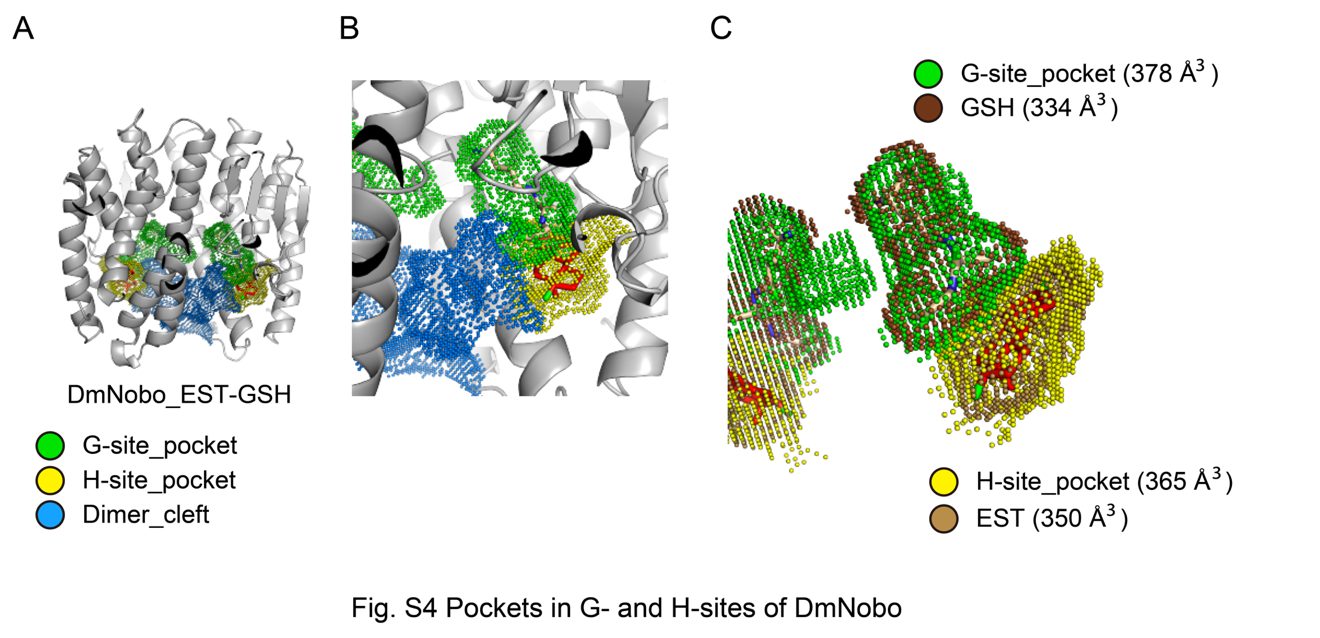

Fig. S4. Pockets in the G- and H-sites of DmNobo

(A, B) Pockets in DmNobo were calculated using 3V. The inner surfaces of the two pockets, i.e., the G- and H-sites, are represented in green and yellow dots, respectively. The cleft between the two subunits of the DmNobo_EST-GSH are shown in blue. The G-site was calculated using the crystal structure of the DmNobo_EST-GSH complex without GSH, and the H-site was calculated using the crystal structure of the DmNobo_EST-GSH complex without EST.

(B) An enlarged view of the G- and H-sites.

(C) The solvent-accessible surfaces of GSH and EST are represented with brown and light brown dots, respectively. The surfaces of the G- and H-sites are superimposed on GSH (green) and EST (yellow).

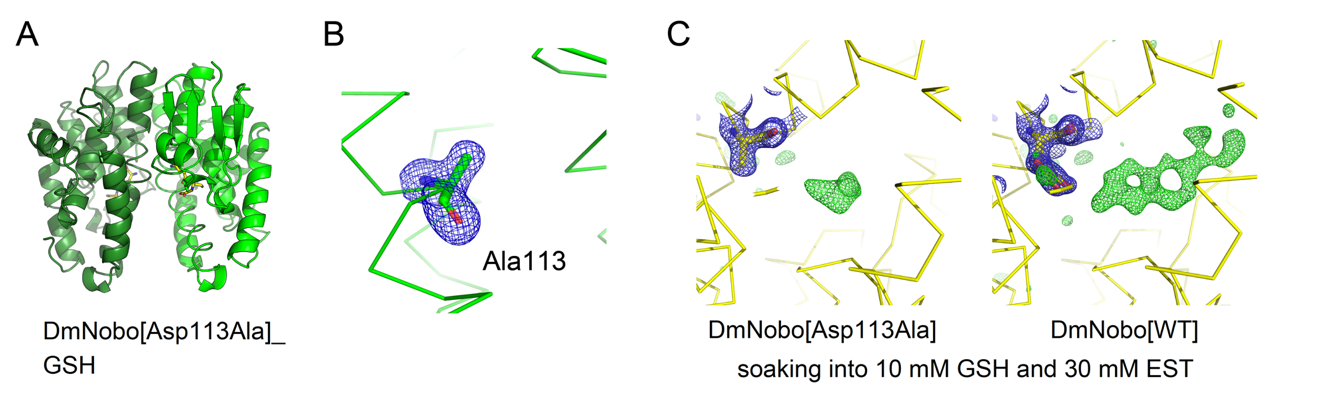

Fig. S5. Characterization of the DmNobo[Asp113Ala] protein.

(A) Overall structure of GSH in complex with DmNobo[Asp113Ala] (DmNobo[Asp113Ala]_GSH). Chains A and B are shaded green and dark green, respectively. The RMSD value from the LSQ fittings between Cα atoms in chain A of DmNobo[WT]_GSH and those of DmNobo[Asp113Ala] was 0.1 Å.

(B) Simulated annealing-omit maps of Ala113 of DmNobo[Asp113Ala]_GSH. m*F*o-D*F*c map (blue) contoured at 4.0σ and within 5 Å from the atoms of Ala113 was overlaid with sticks of Ala113 and a ribbon model of DmNobo[Asp113Ala]_GSH (green).

(C) m*F*o-D*F*c map around the H-sites of DmNobo[Asp113Ala] and DmNobo[WT]. EST did not bind to DmNobo[Asp113Ala]. Crystals of DmNobo[Asp113Ala] and DmNobo[WT] were soaked into an artificial mother liquor (42.5% [w/v] PPG 400 in 100 mM Bis-Tris [pH 6.4]) containing 10 mM GSH and 30 mM EST. m*F*o-D*F*c maps contoured at 4.0σ (green) are shown only within a 5-Å distance from the DmNobo atoms. The blue mesh indicates the 2m*F*o-D*F*c map contoured at 1.5σ around residue 113 (within 5 Å from residue 113).

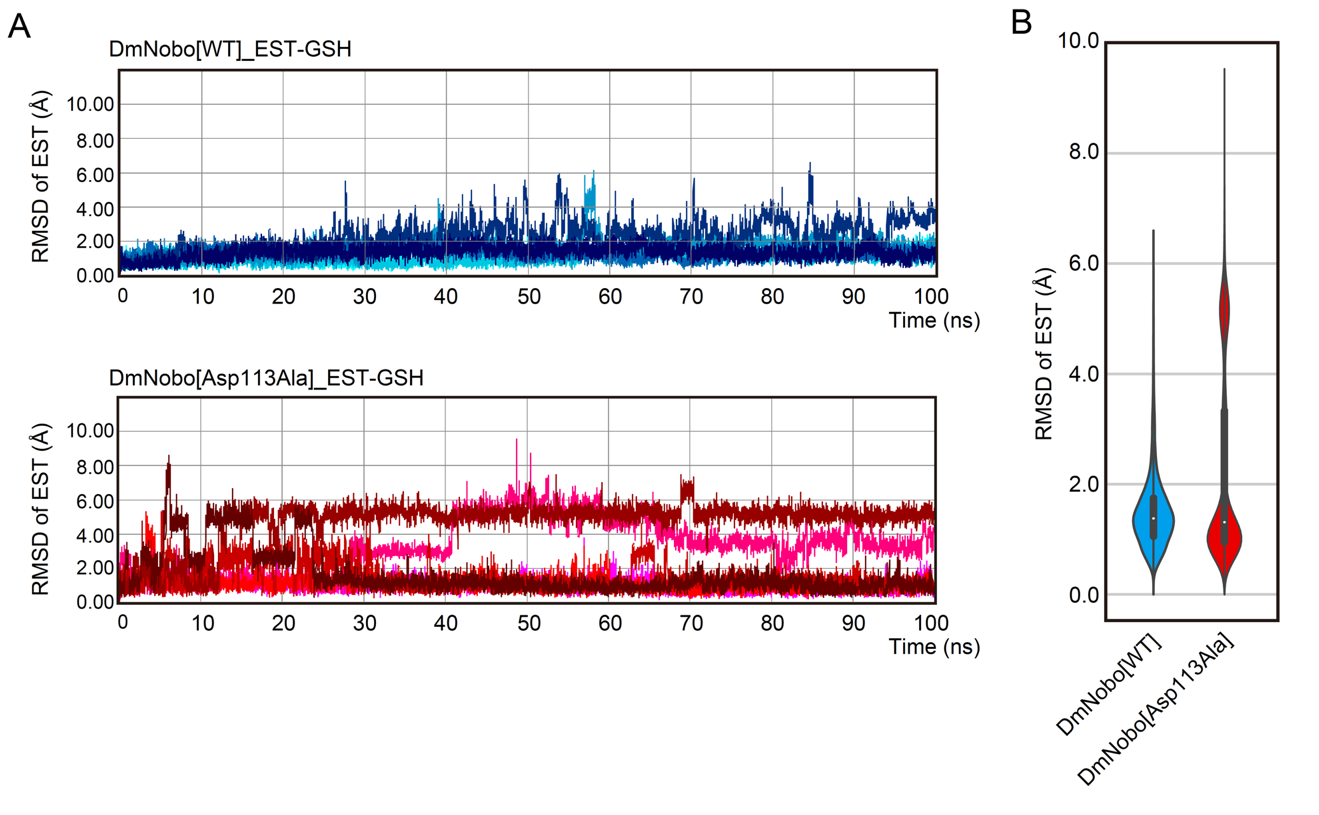

Fig. S6. *In silico* evaluation of the contribution of Asp113 to the interaction between DmNobo and EST.

(A) The RMSD values of EST in DmNobo[WT] (top) or DmNobo[Asp113Ala] (bottom) in triplicate independent calculations. All protein frames were aligned based on the protein backbone of the initial structure. The RMSD (Å) of EST between EST at the initial frame and each subsequent frame is plotted.

(B) Violin-plot of the RMSD of EST in DmNobo[WT] or DmNobo[Asp113Ala]. The median, interquartile range, and 95% confidence interval are shown with a white dot, a bold line, and a narrow line, respectively. The width of the plot indicates the frequency the of frames.

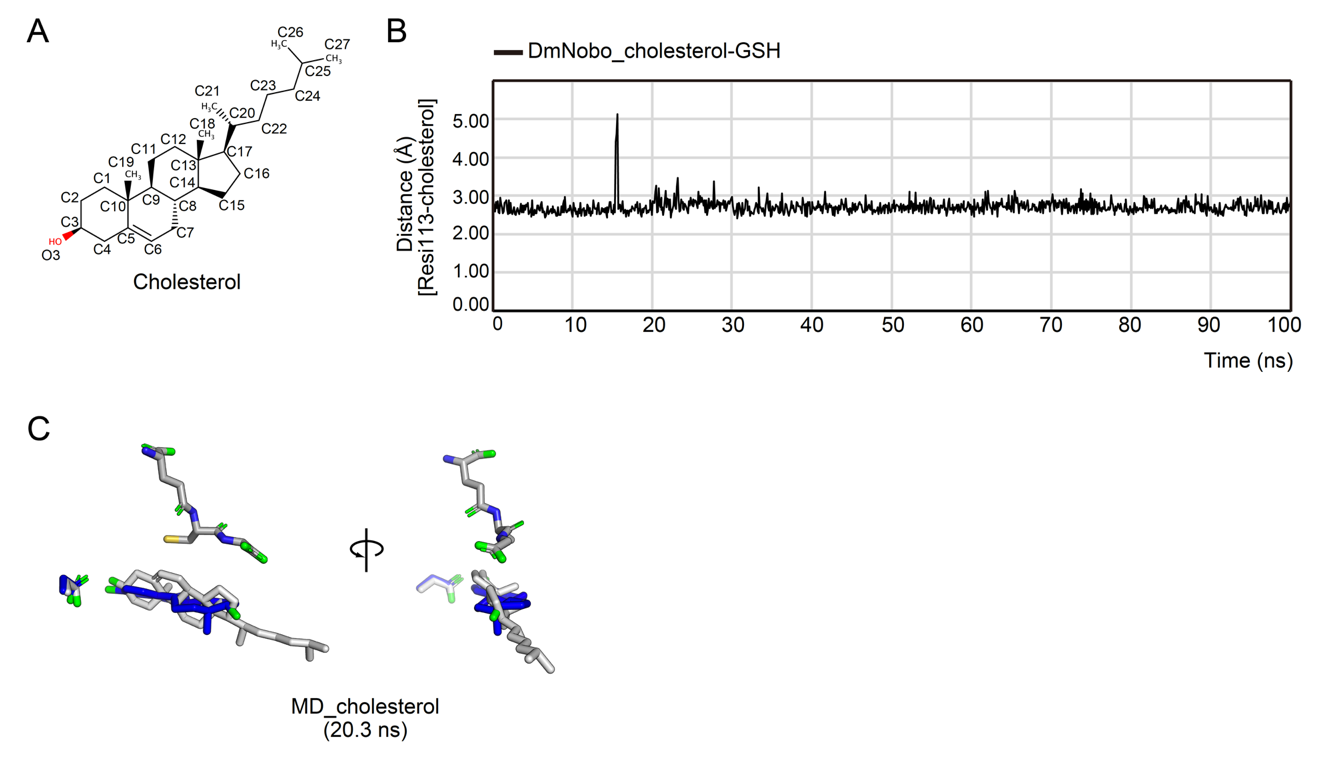

Fig. S7. *In silico* evaluation of interaction between DmNobo and cholesterol

(A) Chemical structure of cholesterol.

(B) MD-simulation results for DmNobo in complex with cholesterol and GSH. The distance between Oδ of Asp113 of DmNobo and O3 of cholesterol was plotted against time.

(C) MD models of DmNobo_cholesterol-GSH and DmNobo_EST-GSH at 20.3 ns.

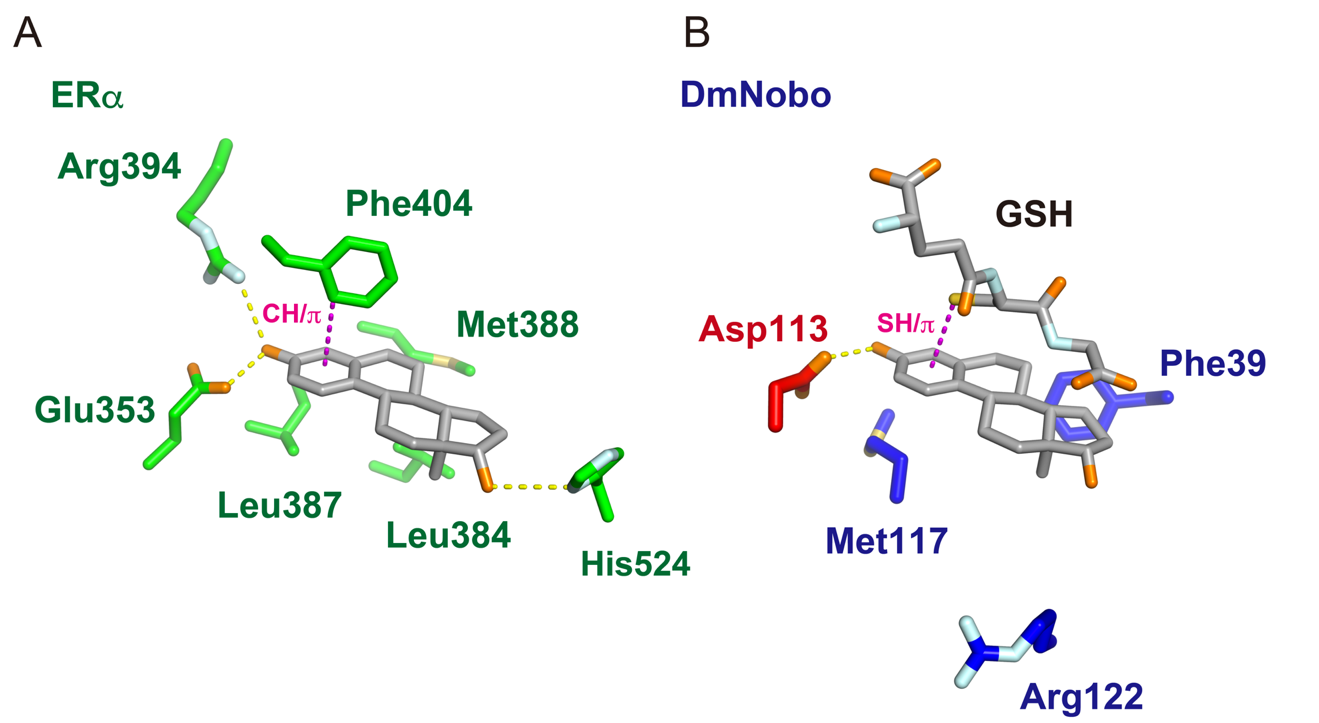

Fig. S8. Comparison of the DmNobo_EST and ERα_EST interactions.

Side chains of representative residues of ERα (A) or DmNobo (B) that interact with EST are represented with sticks (PDB ID for ERα structure = 1QKT). A hydrogen bond between the protein and EST is indicated with a yellow dashed line. A CH/π interaction between Phe404 and EST or an SH/π interaction between GSH and EST is indicated with a pink dashed line.

GATGTTCGCCCTGGGCAGCAGTCTCAATTGGCGCGAGAGGTACATTGTTTAGCGGGCTGGACACAAATCAACTTGATTTTCAGTAATAAAACCAAGAAAATTAAATGTTTTTCTGGTTTAAGTGGGGAAAAATGCAGGTGCTTCTTCTTGTCCAGAAAGTGTGGCCGATCAGCTGAGTCTAACAGTCCGGTTAACAGTGCGTTATAGTATGTCGTTTTCCAGCACTACATCAGGGATACCACACAAATTGAAAAATGAGCGCCAAATTTAAAATGACTTCGTTATCTATCTTATCAATTTCAATTTCCATAGATCTGTACTTTTGGTGAAAGTGAACAGTGTTTATATATTTGATTTTAATATAACAATTAATTGATGAATTATCTCGTAGTATACATTAAAATGCCACCATCAATCAGTACTGTGGCTACAAATGCTTTTCAATTCAGTTACTGATCGACTTTCAAGGCGTTCGTTTGTCTGTTTTTCTACCTGATCCTAACTTAATGGATTACAAGGATCACGATGGCGACTACAAGGACCATGACATCGACTATAAGGACGACGACGATAAGGGATCCTATCCATATGACGTTCCAGATTACGCTATGTCTCAGCCCAAGCCGATTTTGTATTATGATGAGCGCAGCCCACCAGTCCGCAGTTGCCTTATGCTAATCAAATTGCTCGATATAGATGTGGAGCTCAGGTTTGTGAATCTCTTCAAGGGCGAGCAATTCCAAAAAGATTTCTTAGCGGTAAGTAAGTCAAATTTATTCAATGAATAAATTGAATCAATCATTATTTTAAGAAAACGAATAATTTAATTTGCGAAAATTCCTTATAATTATAATTTCTGTTTTATATTTTGCCATTTTTATTTAGTTAAATCCCCAGCACAGTGTTCCCACCCTAGTCCACGGTGATCTGGTGCTGACGGACAGTCATGCTATACTCATTCACCTGGCGGAGAAGTTCGATGAGGGCGGTAGTTTGTGGCCGCAGGAGCACGCAGAACGGATGAAGGTTCTGAACCTCCTGCTCTTCGAGTGCTCCTTTTTGTTCCGACGTGACAGTGATTTTATGGTGGGTGATTCGCAACTAAAACTAGCTGTTTACATATTTTGTTTTTTATTTTTCCCAGTCGGCGACTGGGCGCTCTTCCGCTTCCTCGCTCACTGACTCGCTGCGCTCGGTCGTTCGGCTGCGGCGAGCGGTATCAGCTCACTCAAAGGCGGTAATACGGTTATCCACAGAATCAGGGGATAACGCAGGAAAGAACATGTGAGCAAAAGGCCAGCAAAAGGCCAGGAACCGTAAAAAGGCCGCGTTGCTGGCGTTTTTCCATAGGCTCCGCCCCCCTGACGAGCATCACAAAAATCGACGCTCAAGTCAGAGGTGGCGAAACCCGACAGGACTATAAAGATACCAGGCGTTTCCCCCTGGAAGCTCCCTCGTGCGCTCTCCTGTTCCGACCCTGCCGCTTACCGGATACCTGTCCGCCTTTCTCCCTTCGGGAAGCGTGGCGCTTTCTCATAGCTCACGCTGTAGGTATCTCAGTTCGGTGTAGGTCGTTCGCTCCAAGCTGGGCTGTGTGCACGAACCCCCCGTTCAGCCCGACCGCTGCGCCTTATCCGGTAACTATCGTCTTGAGTCCAACCCGGTAAGACACGACTTATCGCCACTGGCAGCAGCCACTGGTAACAGGATTAGCAGAGCGAGGTATGTAGGCGGTGCTACAGAGTTCTTGAAGTGGTGGCCTAACTACGGCTACACTAGAAGAACAGTATTTGGTATCTGCGCTCTGCTGAAGCCAGTTACCTTCGGAAAAAGAGTTGGTAGCTCTTGATCCGGCAAACAAACCACCGCTGGTAGCGGTGGTTTTTTTGTTTGCAAGCAGCAGATTACGCGCAGAAAAAAAGGATCTCAAGAAGATCCTTTGATCTTTTCTACGGGGTCTGACGCTCAGTGGAACGAAAACTCACGTTAAGGGATTTTGGTCATGAGATTATCAAAAAGGATCTTCACCTAGATCCTTTTAAATTAAAAATGAAGTTTTAAATCAATCTAAAGTATATATGAGTAAACTTGGTCTGACAGTTACCAATGCTTAATCAGTGAGGCACCTATCTCAGCGATCTGTCTATTTCGTTCATCCATAGTTGCCTGACTCCCCGTCGTGTAGATAACTACGATACGGGAGGGCTTACCATCTGGCCCCAGTGCTGCAATGATACCGCGAGATCCACGCTCACCGGCTCCAGATTTATCAGCAATAAACCAGCCAGCCGGAAGGGCCGAGCGCAGAAGTGGTCCTGCAACTTTATCCGCCTCCATCCAGTCTATTAATTGTTGCCGGGAAGCTAGAGTAAGTAGTTCGCCAGTTAATAGTTTGCGCAACGTTGTTGCCATTGCTACAGGCATCGTGGTGTCACGCTCGTCGTTTGGTATGGCTTCATTCAGCTCCGGTTCCCAACGATCAAGGCGAGTTACATGATCCCCCATGTTGTGCAAAAAAGCGGTTAGCTCCTTCGGTCCTCCGATCGTTGTCAGAAGTAAGTTGGCCGCAGTGTTATCACTCATGGTTATGGCAGCACTGCATAATTCTCTTACTGTCATGCCATCCGTAAGATGCTTTTCTGTGACTGGTGAGTACTCAACCAAGTCATTCTGAGAATAGTGTATGCGGCGACCGAGTTGCTCTTGCCCGGCGTCAATACGGGATAATACCGCGCCACATAGCAGAACTTTAAAAGTGCTCATCATTGGAAAACGTTCTTCGGGGCGAAAACTCTCAAGGATCTTACCGCTGTTGAGATCCAGTTCGATGTAACCCACTCGTGCACCCAACTGATCTTCAGCATCTTTTACTTTCACCAGCGTTTCTGGGTGAGCAAAAACAGGAAGGCAAAATGCCGCAAAAAAGGGAATAAGGGCGACACGGAAATGTTGAATACTCATACTCTTCCTTTTTCAATATTATTGAAGCATTTATCAGGGTTATTGTCTCATGAGCGGATACATATTTGAATGTATTTAGAAAAATAAACAAATAGGGGTTCCGCGCACATTTCCCCGAAAAGTGCCACCTGACGTCTAAGAAACCATTATTATCATGACATTAACCTATAAAAATAGGCGTATCACGAGGCCCTTTCGTC

**Fig. S9.**

**DNA sequence of pDonor[KI]-{CG4688_LA}:{3×FLAG/HA/nobo}:{CG4688_RA}**

Gray highlighting: 5′ homology arm; blue highlighting: ATG + 3**×** FLAG tag; yellow highlighting: HA tag; magenta highlighting: exons of Nobo (CG4688); green highlighting: introns of Nobo (CG4688); underlined text: pUC19 ori

GATGTTCGCCCTGGGCAGCAGTCTCAATTGGCGCGAGAGGTACATTGTTTAGCGGGCTGGACACAAATCAACTTGATTTTCAGTAATAAAACCAAGAAAATTAAATGTTTTTCTGGTTTAAGTGGGGAAAAATGCAGGTGCTTCTTCTTGTCCAGAAAGTGTGGCCGATCAGCTGAGTCTAACAGTCCGGTTAACAGTGCGTTATAGTATGTCGTTTTCCAGCACTACATCAGGGATACCACACAAATTGAAAAATGAGCGCCAAATTTAAAATGACTTCGTTATCTATCTTATCAATTTCAATTTCCATAGATCTGTACTTTTGGTGAAAGTGAACAGTGTTTATATATTTGATTTTAATATAACAATTAATTGATGAATTATCTCGTAGTATACATTAAAATGCCACCATCAATCAGTACTGTGGCTACAAATGCTTTTCAATTCAGTTACTGATCGACTTTCAAGGCGTTCGTTTGTCTGTTTTTCTACCTGATCCTAACTTAATGGATTACAAGGATCACGATGGCGACTACAAGGACCATGACATCGACTATAAGGACGACGACGATAAGGGATCCTATCCATATGACGTTCCAGATTACGCTATGTCTCAGCCCAAGCCGATTTTGTATTATGATGAGCGCAGCCCACCAGTCCGCAGTTGCCTTATGCTAATCAAATTGCTCGATATAGATGTGGAGCTCAGGTTTGTGAATCTCTTCAAGGGCGAGCAATTCCAAAAAGATTTCTTAGCGGTAAGTAAGTCAAATTTATTCAATGAATAAATTGAATCAATCATTATTTTAAGAAAACGAATAATTTAATTTGCGAAAATTCCTTATAATTATAATTTCTGTTTTATATTTTGCCATTTTTATTTAGTTAAATCCCCAGCACAGTGTTCCCACCCTAGTCCACGGTGATCTGGTGCTGACGGACAGTCATGCTATACTCATTCACCTGGCGGAGAAGTTCGATGAGGGCGGTAGTTTGTGGCCGCAGGAGCACGCAGAACGGATGAAGGTTCTGAACCTCCTGCTCTTCGAGTGCTCCTTTTTGTTCCGACGT**GCC**AGCGACTTCATGGTGGGTGATTCGCAACTAAAACTAGCTGTTTACATATTTTGTTTTTTATTTTTCCCAGTCGGCGACTGTCCGCCAGGGATTCGCCAATGTCGATGTGGCACATCATGAACGCAAGCTGACCGAGGCGTATATTATCATGGAGCGTTACCTGGAAAATAGCGATTTTATGGCCGGGCCACAGGTAAAAAAAGACCAGCTCATCTGATTGCGATTCCCTGTCGTTGTGGCGGCTCAATTAAGTCTAGAATTTCACCTTTGTTTCCGCGGTTTCAGCTGACGCTCGCCGACTTATCCATCGTGACCACATTGAGCACCGTCAATCTCATGTTTCCCCTGTCGCAGTTCCCACGTCTGCGGCGCTGGTTCACCGCGATGCAGCAGCTGGATGCCTACGAGGCCAACTGCAGTGGCTTGGAGAAGCTCCGCCAAACGATGGAGAGCGTCGGTAGCTTTCAGTTCCCATCGTCATCAGCGGTAGTCACCGAGAAGGTGGAGTAGGGCGCTCTTCCGCTTCCTCGCTCACTGACTCGCTGCGCTCGGTCGTTCGGCTGCGGCGAGCGGTATCAGCTCACTCAAAGGCGGTAATACGGTTATCCACAGAATCAGGGGATAACGCAGGAAAGAACATGTGAGCAAAAGGCCAGCAAAAGGCCAGGAACCGTAAAAAGGCCGCGTTGCTGGCGTTTTTCCATAGGCTCCGCCCCCCTGACGAGCATCACAAAAATCGACGCTCAAGTCAGAGGTGGCGAAACCCGACAGGACTATAAAGATACCAGGCGTTTCCCCCTGGAAGCTCCCTCGTGCGCTCTCCTGTTCCGACCCTGCCGCTTACCGGATACCTGTCCGCCTTTCTCCCTTCGGGAAGCGTGGCGCTTTCTCATAGCTCACGCTGTAGGTATCTCAGTTCGGTGTAGGTCGTTCGCTCCAAGCTGGGCTGTGTGCACGAACCCCCCGTTCAGCCCGACCGCTGCGCCTTATCCGGTAACTATCGTCTTGAGTCCAACCCGGTAAGACACGACTTATCGCCACTGGCAGCAGCCACTGGTAACAGGATTAGCAGAGCGAGGTATGTAGGCGGTGCTACAGAGTTCTTGAAGTGGTGGCCTAACTACGGCTACACTAGAAGAACAGTATTTGGTATCTGCGCTCTGCTGAAGCCAGTTACCTTCGGAAAAAGAGTTGGTAGCTCTTGATCCGGCAAACAAACCACCGCTGGTAGCGGTGGTTTTTTTGTTTGCAAGCAGCAGATTACGCGCAGAAAAAAAGGATCTCAAGAAGATCCTTTGATCTTTTCTACGGGGTCTGACGCTCAGTGGAACGAAAACTCACGTTAAGGGATTTTGGTCATGAGATTATCAAAAAGGATCTTCACCTAGATCCTTTTAAATTAAAAATGAAGTTTTAAATCAATCTAAAGTATATATGAGTAAACTTGGTCTGACAGTTACCAATGCTTAATCAGTGAGGCACCTATCTCAGCGATCTGTCTATTTCGTTCATCCATAGTTGCCTGACTCCCCGTCGTGTAGATAACTACGATACGGGAGGGCTTACCATCTGGCCCCAGTGCTGCAATGATACCGCGAGATCCACGCTCACCGGCTCCAGATTTATCAGCAATAAACCAGCCAGCCGGAAGGGCCGAGCGCAGAAGTGGTCCTGCAACTTTATCCGCCTCCATCCAGTCTATTAATTGTTGCCGGGAAGCTAGAGTAAGTAGTTCGCCAGTTAATAGTTTGCGCAACGTTGTTGCCATTGCTACAGGCATCGTGGTGTCACGCTCGTCGTTTGGTATGGCTTCATTCAGCTCCGGTTCCCAACGATCAAGGCGAGTTACATGATCCCCCATGTTGTGCAAAAAAGCGGTTAGCTCCTTCGGTCCTCCGATCGTTGTCAGAAGTAAGTTGGCCGCAGTGTTATCACTCATGGTTATGGCAGCACTGCATAATTCTCTTACTGTCATGCCATCCGTAAGATGCTTTTCTGTGACTGGTGAGTACTCAACCAAGTCATTCTGAGAATAGTGTATGCGGCGACCGAGTTGCTCTTGCCCGGCGTCAATACGGGATAATACCGCGCCACATAGCAGAACTTTAAAAGTGCTCATCATTGGAAAACGTTCTTCGGGGCGAAAACTCTCAAGGATCTTACCGCTGTTGAGATCCAGTTCGATGTAACCCACTCGTGCACCCAACTGATCTTCAGCATCTTTTACTTTCACCAGCGTTTCTGGGTGAGCAAAAACAGGAAGGCAAAATGCCGCAAAAAAGGGAATAAGGGCGACACGGAAATGTTGAATACTCATACTCTTCCTTTTTCAATATTATTGAAGCATTTATCAGGGTTATTGTCTCATGAGCGGATACATATTTGAATGTATTTAGAAAAATAAACAAATAGGGGTTCCGCGCACATTTCCCCGAAAAGTGCCACCTGACGTCTAAGAAACCATTATTATCATGACATTAACCTATAAAAATAGGCGTATCACGAGGCCCTTTCGTC

**Fig. S10.**

**DNA sequence of pDonor[KI]-{CG4688_LA}:{3×FLAG/HA/nobo*D113A}:{CG4688_RA}**

Gray highlighting: 5′ homology arm; blue highlighting: ATG + 3**×** FLAG tag; yellow highlighting: HA tag; magenta highlighting: exons of Nobo (CG4688); green highlighting: introns of Nobo (CG4688); boxed and bold text: substitution with Ala (GAC → GCC); underlined text: pUC19 ori

Table S1. Crystallographic Summary of DmNobo crystal structures

|  | DmNobo_Apo_1 | DmNobo_Apo_2 | DmNobo_GSH | DmNobo_EST | DmNobo_EST-GSH |
| --- | --- | --- | --- | --- | --- |
| Data Collection | | | | | |
| Space group | *P*2_1_2_1_2_1_ | *P*2_1_2_1_2_1_ | *P*2_1_2_1_2_1_ | *P*2_1_2_1_2_1_ | *P*2_1_2_1_2_1_ |
| Cell Dimensions | | | | | |
| *a*, *b*, *c* (Å) | 58.73, 76.75, 104.82 | 59.12 76.56 106.36 | 58.75, 75.67, 107.84 | 58.40, 75.08, 109.02 | 58.38, 75.06, 108.52 |
| α, β, γ (°) | 90.00, 90.00, 90.00 | 90.00, 90.00, 90.00 | 90.00, 90.00, 90.00 | 90.00, 90.00, 90.00 | 90.00, 90.00, 90.00 |
| Resolution* (Å) | 43.28 - 1.40  (1.45 - 1.40) | 42.83 - 1.50  (1.55 - 1.50) | 32.47 - 1.75  (1.81 - 1.75) | 31.58 - 1.58  (1.64 - 1.58) | 32.59 - 1.55  (1.61 - 1.55) |
| *R*_merge_ | 0.10 (0.66) | 0.05 (0.38) | 0.08 (1.01) | 0.05 (0.84) | 0.04 (0.36) |
| I/σI | 23.2 (3.3) | 33.8 (6.53) | 27.3 (2.3) | 37.6 (2.6) | 42.7 (3.3) |
| Completeness | 0.96 (0.91) | 1.00 (1.00) | 1.00 (0.97) | 0.87 (0.47) | 0.94 (0.72) |
| Redundancy | 13.6 (11.2) | 13.1 (12.7) | 14.3 (12.2) | 14.1 (10.6) | 12.3 (4.1) |
| Refinement | | | | | |
| Resolution (Å) | 43.28 - 1.40 | 42.83 - 1.50 | 32.47 - 1.75 | 31.58 - 1.58 | 32.59 - 1.55 |
| No. reflections | 91,275 | 77,842 | 49,095 | 57,950 | 65,906 |
| *R*_work_ | 0.156 | 0.167 | 0.173 | 0.178 | 0.161 |
| *R*_free_ | 0.180 | 0.193 | 0.207 | 0.209 | 0.182 |
| No. of Atoms | | | | | |
| Protein | 3,670 | 3,880 | 3,609 | 3,700 | 3,817 |
| GSH | 0 | 0 | 40 | 0 | 40 |
| EST | 0 | 0 | 0 | 40 | 40 |
| Water | 536 | 643 | 408 | 408 | 499 |
| *B* factors | | | | | |
| Protein | 16.2 | 22.0 | 22.0 | 24.8 | 20.1 |
| GSH | - | - | 17.7 | - | 15.5 |
| EST | - | - | - | 31.5 | 20.8 |
| Waters | 31.1 | 39.5 | 33.7 | 37.1 | 34.7 |
| RMSD | | | | | |
| Bond length (Å) | 0.013 | 0.008 | 0.007 | 0.011 | 0.010 |
| Bond angles (°) | 1.27 | 0.98 | 0.89 | 1.13 | 1.19 |
| PDB | | | | | |
| ID | 6KEL | 6KEM | 6KEN | 6KEO | 6KEP |

Each structure was determined from diffraction data from one crystal. PDB, Protein Data Bank; RMSD, root-mean-square deviation. *Highest resolution shell is shown in parentheses.

Table S1. Crystallographic Summary of DmNobo crystal structures *Continued*

|  | DmNobo[Asp113Ala]_Apo | DmNobo[Asp113Ala]_GSH |
| --- | --- | --- |
| Data Collection | | |
| Space group | *P*2_1_2_1_2_1_ | *P*2_1_2_1_2_1_ |
| Cell Dimensions | | |
| *a*, *b*, *c* (Å) | 58.72, 75.48, 107.11 | 58.37, 74.83, 107.42 |
| α, β, γ (°) | 90.00, 90.00, 90.00 | 90.00, 90.00, 90.00 |
| Resolution* (Å) | 46.34 - 1.84 (1.90 - 1.84) | 46.03 - 1.84 (1.90 - 1.84) |
| *R*_merge_ | 0.07 (1.10) | 0.11 (1.12) |
| I/σI | 21.3 (2.1) | 15.2 (2.12) |
| Completeness | 1.00 (0.99) | 99.9 (99.5) |
| Redundancy | 13.3 (13.8) | 13.5 (13.9) |
| Refinement | | |
| Resolution (Å) | 46.34 - 1.84 | 46.03 - 1.84 |
| No. reflections | 42,099 | 41,768 |
| *R*_work_ | 0.209 | 0.189 |
| *R*_free_ | 0.241 | 0.221 |
| No. of Atoms | | |
| Protein | 3,495 | 3,529 |
| GSH | 0 | 40 |
| EST | 0 | 0 |
| Water | 174 | 146 |
| *B* factors | | |
| Protein | 33.45 | 21.9 |
| GSH | - | 18.1 |
| EST | - | - |
| Waters | 36.71 | 26.5 |
| RMSD | | |
| Bond length (Å) | 0.005 | 0.004 |
| Bond angles (°) | 0.73 | 0.66 |
| PDB | | |
| ID | 6KEQ | 6KER |

**Table S2. root-mean-square deviation (RMSD) among DmNobo crystal structures**

|  |  | RMSD (Å) of Cα atoms of chain A^†^/  RMSD (Å) of Cα atoms of chains A and B^‡^ | | |
| --- | --- | --- | --- | --- |
|  | Coordinate error (Å)^*^ | DmNobo_GSH | DmNobo_EST | DmNobo_EST-GSH |
| DmNobo_Apo_2 | 0.13 | 0.20 / 0.35 | 0.25 / 0.42 | 0.48 / 0.55 |
| DmNobo_GSH | 0.20 |  | 0.16 / 0.24 | 0.42 / 0.36 |
| DmNobo_EST | 0.19 |  |  | 0.41 / 0.33 |
| DmNobo_EST-GSH | 0.15 |  |  |  |

^*^ Coordinate errors are estimated by a maximum-likelihood method.

^†^ Number of aligned Cα atoms in chain A: 198 atoms

^‡^ Number of aligned Cα atoms in chain A and B: 388 atoms

**Table S3. Summary of H-site-composing, or EST-interacting atoms in DmNobo**

| Residue | H-site-  composing  atoms | EST-interacting atoms^*^ | DmNobo-interacting atoms^†^ | Distance ^‡^  (Å) | Identity  among  DmGST  D/E/T | Identity  among  Nobo | Total IFIE  to EST  (kcal/mol) |
| --- | --- | --- | --- | --- | --- | --- | --- |
| Arg13 | C, Cβ | - | - | - | 0.07 | 0.05 | -0.31 |
| Ser14 | Cα, N, Cβ | - | - | - | 0.85 | 1.00 | -1.78 |
| Pro15 | Cδ, Cγ | Cδ | O3 | 3.2 | 0.62 | 1.00 | -3.93 |
|  |  | Cδ | C3 | 3.6 |  |  |  |
|  |  | Cδ | C4 | 3.7 |  |  |  |
|  |  | Cγ | O3 | 3.1 |  |  |  |
| Leu38 | C, O, Cβ, Cδ | Cβ | C15 | 4.0 | 0.61 | 1.00 | -1.86 |
|  |  | Cδ | C7 | 4.0 |  |  |  |
| Phe39 | Cα, Cδ, Cγ, Cε, Cζ | Cε1 | C6 | 3.9 | 0.07 | 1.00 | -6.77 |
|  |  |  | C7 | 3.7 |  |  |  |
|  |  | π | C15 | 3.9 |  |  |  |
| Gln43 | Nε | - | - | - | 0.21 | 0.05 | -0.51 |
| Phe110 | Cε, Cζ | Cε | C2 | 4.0 | 0.42 | 1.00 | -2.79 |
| Asp113 | Oδ | Oδ | O3 | 2.6 | 0.01 | 0.95 | -41.4 |
| Ser114 | Cα, Cβ | Cα | C1 | 3.8 | 0.10 | 0.76 | -2.17 |
|  |  | Cα | C2 | 3.8 |  |  |  |
|  |  | Cβ | C1 | 3.9 |  |  |  |
| Met117 | Cβ, Cγ, Cε | Cβ | C1 | 3.9 | 0.04 | 0.40 | -3.52 |
|  |  | Cβ | C2 | 3.9 |  |  |  |
|  |  | Cγ | C2 | 3.8 |  |  |  |
|  |  | Cγ | C3 | 3.9 |  |  |  |
| Ser118 | Cα, N, Cβ, Cγ | - | - | - | 0.06 | 0.33 | -3.25 |
| Val121 | Cβ, Cγ1, Cγ2 | Cβ | C18 | 3.9 | 0.11 | 0.05 | -1.63 |
|  |  | Cγ1 | C18 | 3.9 |  |  |  |
|  |  | Cγ2 | C18 | 3.8 |  |  |  |
| Arg122 | Cβ | - | - | - | 0.07 | 0.14 | -1.49 |
| Thr172 | Cγ | Cγ | O3 | 3.3 | 0.31 | 0.52 | 0.026 |
| Leu208 | Cδ1, Cδ2 | Cδ1 | C4 | 3.9 | 0.31 | 0.95 | -2.17 |
|  |  | Cδ1 | C6 | 3.8 |  |  |  |
|  |  | Cδ2 | O3 | 4.0 |  |  |  |
|  |  | Cδ2 | C4 | 3.9 |  |  |  |
| Met212 | Sδ | - | - | - | 0.05 | 0.95 | -0.65 |
| GSH | Cys_Cβ, Cys_Sγ | Cys_Sγ | π | 3.7 | - | - | -3.48 |

^*^ Atoms of DmNobo within 4.0 Å from EST.

^†^ Atoms of EST within 4.0 Å from each EST-interacting atom.

^‡^ Average of chains A and B in the asymmetric unit.

**Table S4. FMO analysis between fragments in DmNobo and EST**

| Residue | Total | ES | EX | CT | DI |
| --- | --- | --- | --- | --- | --- |
|  | [kcal/mol] | [kcal/mol] | [kcal/mol] | [kcal/mol] | [kcal/mol] |
| Arg13 | -0.31 | 0.36 | 0.03 | -0.10 | -0.59 |
| Ser14 | -1.78 | 0.49 | 0.20 | -1.01 | -1.47 |
| Pro15 | -3.93 | -1.52 | 2.53 | -1.25 | -3.69 |
| Arg18 | -1.85 | -1.66 | 0.00 | -0.02 | -0.17 |
| Leu38 | -1.86 | -0.67 | 2.75 | -0.79 | -3.15 |
| Phe39 | -6.77 | -1.77 | 4.34 | -1.97 | -7.37 |
| Lys40 | 0.13 | 0.28 | 0.00 | 0.00 | -0.15 |
| Gln43 | -0.51 | 0.25 | 0.12 | -0.18 | -0.69 |
| Phe110 | -2.79 | -1.08 | 0.31 | -0.51 | -1.51 |
| Asp113 | -41.38 | -49.71 | 26.87 | -12.63 | -5.91 |
| Ser114 | -2.17 | -0.03 | 3.72 | -2.76 | -3.11 |
| Asp115 | 3.31 | 1.66 | 0.07 | 2.18 | -0.59 |
| Met117 | -3.52 | 0.12 | 2.12 | -1.36 | -4.39 |
| Ser118 | -3.25 | -0.41 | 0.87 | -1.05 | -2.65 |
| Ala119 | -0.36 | -0.24 | 0.03 | 0.24 | -0.39 |
| Val121 | -1.63 | -0.74 | 2.35 | -0.47 | -2.77 |
| Arg122 | -1.49 | -1.10 | 0.00 | -0.03 | -0.36 |
| Leu208 | -2.17 | -0.74 | 2.23 | -0.70 | -2.97 |
| GSH_Gly | -2.49 | 0.64 | 0.32 | -0.89 | -2.56 |
| GSH_Cys | -3.48 | -1.00 | 5.12 | -1.68 | -5.92 |
| PIEDA | -82.4 | -59.2 | 55.8 | -25.8 | -53.1 |

**Table S5. ID of amino acid sequences used in phylogenetic analysis**

| Accession | Description |
| --- | --- |
| Query_10001 | tr\|C4WSG3\|C4WSG3_ACYPI ACYPI009122 protein OS=Acyrthosiphon pisum OX=7029 GN=ACYPI009122 PE=2 SV=1 |
| Query_10002 | tr\|C4WT83\|C4WT83_ACYPI ACYPI008657 protein OS=Acyrthosiphon pisum OX=7029 GN=ACYPI008657 PE=2 SV=1 |
| Query_10003 | tr\|C4WUF9\|C4WUF9_ACYPI ACYPI005620 protein OS=Acyrthosiphon pisum OX=7029 GN=ACYPI005620 PE=2 SV=1 |
| Query_10004 | tr\|I6SCG3\|I6SCG3_ACYPI Glutathione S-transferase D5 OS=Acyrthosiphon pisum OX=7029 GN=GstD5 PE=3 SV=1 |
| Query_10005 | tr\|J9JLJ3\|J9JLJ3_ACYPI Uncharacterized protein OS=Acyrthosiphon pisum OX=7029 PE=4 SV=2 |
| Query_10006 | tr\|J9K2A8\|J9K2A8_ACYPI Uncharacterized protein OS=Acyrthosiphon pisum OX=7029 PE=3 SV=1 |
| Query_10007 | tr\|J9K372\|J9K372_ACYPI Uncharacterized protein OS=Acyrthosiphon pisum OX=7029 PE=3 SV=1 |
| Query_10008 | tr\|J9K6N7\|J9K6N7_ACYPI Uncharacterized protein OS=Acyrthosiphon pisum OX=7029 GN=100167788 PE=4 SV=1 |
| Query_10009 | tr\|J9M2I6\|J9M2I6_ACYPI Uncharacterized protein OS=Acyrthosiphon pisum OX=7029 PE=3 SV=2 |
| Query_10010 | tr\|X1WJM1\|X1WJM1_ACYPI Uncharacterized protein OS=Acyrthosiphon pisum OX=7029 PE=3 SV=1 |
| Query_10011 | tr\|Q17MA9\|Q17MA9_AEDAE AAEL001061-PA OS=Aedes aegypti OX=7159 GN=GSTD1 PE=4 SV=1 |
| Query_10012 | tr\|Q174V0\|Q174V0_AEDAE AAEL006764-PA (Fragment) OS=Aedes aegypti OX=7159 GN=AAEL006764 PE=4 SV=1 |
| Query_10013 | tr\|Q17MB0\|Q17MB0_AEDAE AAEL001078-PA OS=Aedes aegypti OX=7159 GN=GSTD2 PE=4 SV=1 |
| Query_10014 | tr\|Q17MB7\|Q17MB7_AEDAE AAEL001059-PA OS=Aedes aegypti OX=7159 GN=GSTD3 PE=3 SV=1 |
| Query_10015 | tr\|Q17MB8\|Q17MB8_AEDAE AAEL001054-PA (Fragment) OS=Aedes aegypti OX=7159 GN=GSTD4 PE=4 SV=2 |
| Query_10016 | tr\|Q17MB1\|Q17MB1_AEDAE AAEL001071-PA OS=Aedes aegypti OX=7159 GN=GSTD5 PE=4 SV=1 |
| Query_10017 | tr\|Q16SH6\|Q16SH6_AEDAE AAEL010591-PA OS=Aedes aegypti OX=7159 GN=GSTD6 PE=3 SV=1 |
| Query_10018 | tr\|Q17MA8\|Q17MA8_AEDAE AAEL001090-PA OS=Aedes aegypti OX=7159 GN=GSTD7 PE=4 SV=1 |
| Query_10019 | tr\|Q170D2\|Q170D2_AEDAE AAEL007954-PA OS=Aedes aegypti OX=7159 GN=GSTE1 PE=3 SV=2 |
| Query_10020 | tr\|Q5PY77\|Q5PY77_AEDAE AAEL007951-PA OS=Aedes aegypti OX=7159 GN=GSTe2 PE=1 SV=1 |
| Query_10021 | tr\|Q170C6\|Q170C6_AEDAE AAEL007947-PA OS=Aedes aegypti OX=7159 GN=GSTE3 PE=3 SV=1 |
| Query_10022 | tr\|Q5PY78\|Q5PY78_AEDAE AAEL007962-PA OS=Aedes aegypti OX=7159 GN=GSTe4 PE=2 SV=1 |
| Query_10023 | tr\|Q170C9\|Q170C9_AEDAE AAEL007964-PA OS=Aedes aegypti OX=7159 GN=GSTE5 PE=3 SV=1 |
| Query_10024 | tr\|Q170C7\|Q170C7_AEDAE AAEL007946-PA OS=Aedes aegypti OX=7159 GN=GSTE6 PE=4 SV=2 |
| Query_10025 | tr\|Q170C8\|Q170C8_AEDAE AAEL007948-PA OS=Aedes aegypti OX=7159 GN=GSTE7 PE=3 SV=1 |
| Query_10026 | tr\|Q170D3\|Q170D3_AEDAE AAEL007955-PA (Fragment) OS=Aedes aegypti OX=7159 GN=GSTE8 PE=4 SV=1 |
| Query_10027 | tr\|Q5PY76\|Q5PY76_AEDAE AAEL009017-PA (Fragment) OS=Aedes aegypti OX=7159 GN=GSTt1 PE=2 SV=1 |
| Query_10028 | tr\|Q16X19\|Q16X19_AEDAE AAEL009016-PA OS=Aedes aegypti OX=7159 GN=GSTT2 PE=3 SV=2 |
| Query_10029 | tr\|Q16X21\|Q16X21_AEDAE AAEL009020-PA OS=Aedes aegypti OX=7159 GN=GSTT3 PE=3 SV=2 |
| Query_10030 | tr\|Q17DF6\|Q17DF6_AEDAE AAEL004229-PA OS=Aedes aegypti OX=7159 GN=GSTT4 PE=3 SV=1 |
| Query_10031 | tr\|A0A023ENG1\|A0A023ENG1_AEDAL Putative cpij002683 glutathione s-transferase 1-1 OS=Aedes albopictus OX=7160 GN=RP20_CCG018982 PE=2 SV=1 |
| Query_10032 | tr\|A0A023EKA0\|A0A023EKA0_AEDAL Uncharacterized protein OS=Aedes albopictus OX=7160 GN=109402247 PE=2 SV=1 |
| Query_10033 | tr\|A0A182GCK7\|A0A182GCK7_AEDAL Uncharacterized protein OS=Aedes albopictus OX=7160 GN=RP20_CCG007573 PE=3 SV=1 |
| Query_10034 | tr\|A0A182GEB4\|A0A182GEB4_AEDAL Uncharacterized protein OS=Aedes albopictus OX=7160 GN=RP20_CCG008439 PE=3 SV=1 |
| Query_10035 | tr\|A0A182GVH4\|A0A182GVH4_AEDAL Uncharacterized protein OS=Aedes albopictus OX=7160 GN=RP20_CCG016463 PE=3 SV=1 |
| Query_10036 | tr\|A0A182GCK8\|A0A182GCK8_AEDAL Uncharacterized protein OS=Aedes albopictus OX=7160 GN=109424803 PE=3 SV=1 |
| Query_10037 | tr\|A0A023EK27\|A0A023EK27_AEDAL Putative glutathione s-transferase OS=Aedes albopictus OX=7160 PE=2 SV=1 |
| Query_10038 | tr\|A0A023EMF5\|A0A023EMF5_AEDAL Putative glutathione s-transferase e4 OS=Aedes albopictus OX=7160 PE=2 SV=1 |
| Query_10039 | tr\|A0A023EK11\|A0A023EK11_AEDAL Putative glutathione s-transferase e2 OS=Aedes albopictus OX=7160 PE=2 SV=1 |
| Query_10040 | tr\|A0A023EK53\|A0A023EK53_AEDAL Putative glutathione s-transferase e4 OS=Aedes albopictus OX=7160 PE=2 SV=1 |
| Query_10041 | tr\|A0A1W7R8Y6\|A0A1W7R8Y6_AEDAL Putative glutathione s-transferase OS=Aedes albopictus OX=7160 PE=3 SV=1 |
| Query_10042 | tr\|A0A023EJP7\|A0A023EJP7_AEDAL Putative glutathione s-transferase 1 isoform d OS=Aedes albopictus OX=7160 PE=2 SV=1 |
| Query_10043 | tr\|A0A182GJY7\|A0A182GJY7_AEDAL Uncharacterized protein OS=Aedes albopictus OX=7160 GN=RP20_CCG011233 PE=3 SV=1 |
| Query_10044 | tr\|A0A182GCK6\|A0A182GCK6_AEDAL Uncharacterized protein OS=Aedes albopictus OX=7160 GN=RP20_CCG007572 PE=3 SV=1 |
| Query_10045 | tr\|A0A182GEB7\|A0A182GEB7_AEDAL Uncharacterized protein OS=Aedes albopictus OX=7160 GN=RP20_CCG008442 PE=3 SV=1 |
| Query_10046 | tr\|A0A182H003\|A0A182H003_AEDAL Uncharacterized protein OS=Aedes albopictus OX=7160 GN=RP20_CCG018985 PE=3 SV=1 |
| Query_10047 | tr\|A0A182GYD0\|A0A182GYD0_AEDAL Uncharacterized protein OS=Aedes albopictus OX=7160 GN=RP20_CCG018089 PE=3 SV=1 |
| Query_10048 | tr\|A0A182GHZ1\|A0A182GHZ1_AEDAL Uncharacterized protein OS=Aedes albopictus OX=7160 GN=RP20_CCG010237 PE=3 SV=1 |
| Query_10049 | tr\|A0A182GYD1\|A0A182GYD1_AEDAL Uncharacterized protein OS=Aedes albopictus OX=7160 GN=RP20_CCG018090 PE=3 SV=1 |
| Query_10050 | tr\|A0A182GYC8\|A0A182GYC8_AEDAL Uncharacterized protein OS=Aedes albopictus OX=7160 GN=RP20_CCG018087 PE=3 SV=1 |
| Query_10051 | tr\|A0A182GYC5\|A0A182GYC5_AEDAL Uncharacterized protein OS=Aedes albopictus OX=7160 GN=RP20_CCG018084 PE=3 SV=1 |
| Query_10052 | tr\|A0A182H001\|A0A182H001_AEDAL Uncharacterized protein OS=Aedes albopictus OX=7160 GN=RP20_CCG018983 PE=3 SV=1 |
| Query_10053 | tr\|A0A182GYC6\|A0A182GYC6_AEDAL Uncharacterized protein OS=Aedes albopictus OX=7160 GN=RP20_CCG018085 PE=3 SV=1 |
| Query_10054 | tr\|A0A182G345\|A0A182G345_AEDAL Uncharacterized protein OS=Aedes albopictus OX=7160 GN=RP20_CCG001852 PE=3 SV=1 |
| Query_10055 | tr\|A0A182H002\|A0A182H002_AEDAL Uncharacterized protein OS=Aedes albopictus OX=7160 GN=RP20_CCG018984 PE=3 SV=1 |
| Query_10056 | tr\|A0A1W7R8X7\|A0A1W7R8X7_AEDAL Putative glutathione s-transferase e4 (Fragment) OS=Aedes albopictus OX=7160 PE=4 SV=1 |
| Query_10057 | tr\|A0A182GEB5\|A0A182GEB5_AEDAL Uncharacterized protein OS=Aedes albopictus OX=7160 GN=RP20_CCG008440 PE=4 SV=1 |
| Query_10058 | tr\|A0A182GYC7\|A0A182GYC7_AEDAL Uncharacterized protein OS=Aedes albopictus OX=7160 GN=RP20_CCG018086 PE=4 SV=1 |
| Query_10059 | tr\|A0A182GYC9\|A0A182GYC9_AEDAL Uncharacterized protein OS=Aedes albopictus OX=7160 GN=RP20_CCG018088 PE=4 SV=1 |
| Query_10060 | tr\|A0A182GEB6\|A0A182GEB6_AEDAL Uncharacterized protein OS=Aedes albopictus OX=7160 PE=4 SV=1 |
| Query_10061 | XP_013192421.1 PREDICTED: glutathione S-transferase 1-like [Amyelois transitella] |
| Query_10062 | XP_013189941.1 PREDICTED: glutathione S-transferase 1-like [Amyelois transitella] |
| Query_10063 | XP_013196516.1 PREDICTED: glutathione S-transferase 1-like [Amyelois transitella] |
| Query_10064 | XP_013187010.1 PREDICTED: glutathione S-transferase 1-like [Amyelois transitella] |
| Query_10065 | XP_013196509.1 PREDICTED: glutathione S-transferase 1, isoform C-like [Amyelois transitella] |
| Query_10066 | XP_013194108.1 PREDICTED: uncharacterized protein LOC106137747 [Amyelois transitella] |
| Query_10067 | XP_013196508.1 PREDICTED: glutathione S-transferase 1-1-like [Amyelois transitella] |
| Query_10068 | XP_013198047.1 PREDICTED: glutathione S-transferase 1-like [Amyelois transitella] |
| Query_10069 | XP_013187696.1 PREDICTED: glutathione S-transferase 1, isoform C-like [Amyelois transitella] |
| Query_10070 | XP_013182960.1 PREDICTED: glutathione S-transferase 1-like [Amyelois transitella] |
| Query_10071 | NP_001299595.1 uncharacterized protein LOC106136399 precursor [Amyelois transitella] |
| Query_10072 | XP_013187697.1 PREDICTED: glutathione S-transferase 1, isoform C-like [Amyelois transitella] |
| Query_10073 | XP_013192447.1 PREDICTED: glutathione S-transferase 1-like [Amyelois transitella] |
| Query_10074 | XP_013194098.1 PREDICTED: glutathione S-transferase 1-like [Amyelois transitella] |
| Query_10075 | tr\|W5JW71\|W5JW71_ANODA Glutathione transferase, delta class OS=Anopheles darlingi OX=43151 GN=AND_000801 PE=3 SV=1 |
| Query_10076 | tr\|W5JXC1\|W5JXC1_ANODA Glutathione transferase, theta class OS=Anopheles darlingi OX=43151 GN=AND_000084 PE=3 SV=1 |
| Query_10077 | tr\|W5JWL4\|W5JWL4_ANODA Glutathione transferase, theta class OS=Anopheles darlingi OX=43151 GN=AND_000085 PE=3 SV=1 |
| Query_10078 | tr\|W5JWI4\|W5JWI4_ANODA Glutathione S-transferase 1-6 OS=Anopheles darlingi OX=43151 GN=AND_000834 PE=3 SV=1 |
| Query_10079 | tr\|W5J8C0\|W5J8C0_ANODA Glutathione transferase OS=Anopheles darlingi OX=43151 GN=AND_008209 PE=3 SV=1 |
| Query_10080 | tr\|W5J9X2\|W5J9X2_ANODA Glutathione S-transferase, epsilon class OS=Anopheles darlingi OX=43151 GN=AND_008205 PE=3 SV=1 |
| Query_10081 | tr\|W5JTD5\|W5JTD5_ANODA Glutathione transferase OS=Anopheles darlingi OX=43151 GN=AND_000803 PE=3 SV=1 |
| Query_10082 | tr\|W5J8C4\|W5J8C4_ANODA Glutathione S-transferase, epsilon class OS=Anopheles darlingi OX=43151 GN=AND_008208 PE=3 SV=1 |
| Query_10083 | tr\|W5JA16\|W5JA16_ANODA Glutathione S-transferase theta-2 OS=Anopheles darlingi OX=43151 GN=AND_008797 PE=3 SV=1 |
| Query_10084 | tr\|W5J546\|W5J546_ANODA Glutathione transferase OS=Anopheles darlingi OX=43151 GN=AND_008798 PE=3 SV=1 |
| Query_10085 | tr\|W5JJP4\|W5JJP4_ANODA Glutathione transferase, delta class OS=Anopheles darlingi OX=43151 GN=AND_004770 PE=3 SV=1 |
| Query_10086 | tr\|W5JBQ5\|W5JBQ5_ANODA Glutathione S-transferase, epsilon class OS=Anopheles darlingi OX=43151 GN=AND_008212 PE=3 SV=1 |
| Query_10087 | tr\|W5JVI0\|W5JVI0_ANODA Glutathione S transferase D1 OS=Anopheles darlingi OX=43151 GN=AND_000833 PE=3 SV=1 |
| Query_10088 | tr\|W5J6X4\|W5J6X4_ANODA Glutathione S-transferase, epsilon class OS=Anopheles darlingi OX=43151 GN=AND_008200 PE=3 SV=1 |
| Query_10089 | tr\|W5JSQ1\|W5JSQ1_ANODA Glutathione transferase OS=Anopheles darlingi OX=43151 GN=AND_000804 PE=4 SV=1 |
| Query_10090 | tr\|A0A2M4CU45\|A0A2M4CU45_ANODA Putative chain a glutathione s-transferase OS=Anopheles darlingi OX=43151 PE=3 SV=1 |
| Query_10091 | tr\|A0A2M4CV43\|A0A2M4CV43_ANODA Putative chain a glutathione s-transferase OS=Anopheles darlingi OX=43151 PE=3 SV=1 |
| Query_10092 | tr\|A0A2M4CJX8\|A0A2M4CJX8_ANODA Putative glutathione s-transferase 1 isoform d (Fragment) OS=Anopheles darlingi OX=43151 PE=3 SV=1 |
| Query_10093 | tr\|A0A2M4CYI6\|A0A2M4CYI6_ANODA Putative glutathione s-transferase epsilon class OS=Anopheles darlingi OX=43151 PE=3 SV=1 |
| Query_10094 | tr\|W5JWK8\|W5JWK8_ANODA Glutathione transferase OS=Anopheles darlingi OX=43151 GN=AND_000802 PE=4 SV=1 |
| Query_10095 | tr\|W5J882\|W5J882_ANODA Glutathione transferase OS=Anopheles darlingi OX=43151 GN=AND_008796 PE=4 SV=1 |
| Query_10096 | tr\|W5JBI5\|W5JBI5_ANODA Glutathione transferase OS=Anopheles darlingi OX=43151 GN=AND_008211 PE=4 SV=1 |
| Query_10097 | tr\|W5J6X9\|W5J6X9_ANODA Glutathione S-transferase, epsilon class OS=Anopheles darlingi OX=43151 GN=AND_008204 PE=4 SV=1 |
| Query_10098 | tr\|W5JW53\|W5JW53_ANODA Glutathione transferase, delta class OS=Anopheles darlingi OX=43151 GN=AND_000835 PE=4 SV=1 |
| Query_10099 | tr\|W5JC52\|W5JC52_ANODA Glutathione transferase, delta class OS=Anopheles darlingi OX=43151 GN=AND_007981 PE=4 SV=1 |
| Query_10100 | tr\|A0A084WTH5\|A0A084WTH5_ANOSI Glutathione S-transferase D6 OS=Anopheles sinensis OX=74873 GN=ZHAS_00021747 PE=3 SV=1 |
| Query_10101 | tr\|A0A084VMZ2\|A0A084VMZ2_ANOSI Glutathione s-transferase E5 OS=Anopheles sinensis OX=74873 GN=ZHAS_00006696 PE=3 SV=1 |
| Query_10102 | tr\|A0A084WQQ1\|A0A084WQQ1_ANOSI Glutathione transferase epsilon3 OS=Anopheles sinensis OX=74873 GN=ZHAS_00020795 PE=3 SV=1 |
| Query_10103 | tr\|A0A084WQQ3\|A0A084WQQ3_ANOSI Glutathione s-transferase E2 OS=Anopheles sinensis OX=74873 GN=ZHAS_00020699 PE=3 SV=1 |
| Query_10104 | tr\|A0A084VQI2\|A0A084VQI2_ANOSI AGAP000888-PA-like protein OS=Anopheles sinensis OX=74873 GN=ZHAS_00007603 PE=3 SV=1 |
| Query_10105 | tr\|A0A084VQI3\|A0A084VQI3_ANOSI Uncharacterized protein OS=Anopheles sinensis OX=74873 GN=ZHAS_00007604 PE=3 SV=1 |
| Query_10106 | tr\|A0A084WTH9\|A0A084WTH9_ANOSI AGAP004383-PA-like protein OS=Anopheles sinensis OX=74873 GN=ZHAS_00021752 PE=3 SV=1 |
| Query_10107 | tr\|A0A084WTI1\|A0A084WTI1_ANOSI Uncharacterized protein OS=Anopheles sinensis OX=74873 GN=ZHAS_00021754 PE=3 SV=1 |
| Query_10108 | tr\|A0A084WTH8\|A0A084WTH8_ANOSI AGAP004382-PA-like protein OS=Anopheles sinensis OX=74873 GN=ZHAS_00021751 PE=3 SV=1 |
| Query_10109 | tr\|A0A084WTH6\|A0A084WTH6_ANOSI AGAP004380-PA-like protein OS=Anopheles sinensis OX=74873 GN=ZHAS_00021748 PE=3 SV=1 |
| Query_10110 | tr\|A0A084WTH7\|A0A084WTH7_ANOSI AGAP004382-PA-like protein OS=Anopheles sinensis OX=74873 GN=ZHAS_00021749 PE=3 SV=1 |
| Query_10111 | tr\|A0A084WTI2\|A0A084WTI2_ANOSI AGAP004171-PA-like protein OS=Anopheles sinensis OX=74873 GN=ZHAS_00021755 PE=3 SV=1 |
| Query_10112 | tr\|A0A084WTH4\|A0A084WTH4_ANOSI AGAP004378-PA-like protein OS=Anopheles sinensis OX=74873 GN=ZHAS_00021746 PE=3 SV=1 |
| Query_10113 | tr\|A0A084VUG1\|A0A084VUG1_ANOSI AGAP003257-PA-like protein OS=Anopheles sinensis OX=74873 GN=ZHAS_00009226 PE=3 SV=1 |
| Query_10114 | tr\|A0A084VMZ0\|A0A084VMZ0_ANOSI AGAP009190-PA-like protein OS=Anopheles sinensis OX=74873 GN=ZHAS_00006694 PE=3 SV=1 |
| Query_10115 | tr\|A0A084WTJ6\|A0A084WTJ6_ANOSI AGAP004164-PB-like protein OS=Anopheles sinensis OX=74873 GN=ZHAS_00021770 PE=3 SV=1 |
| Query_10116 | tr\|A0A084VMZ4\|A0A084VMZ4_ANOSI Glutathione s-transferase E2 OS=Anopheles sinensis OX=74873 GN=ZHAS_00006698 PE=4 SV=1 |
| Query_10117 | tr\|A0A084VMZ3\|A0A084VMZ3_ANOSI Glutathione s-transferase E4 OS=Anopheles sinensis OX=74873 GN=ZHAS_00006697 PE=4 SV=1 |
| Query_10118 | tr\|A0A084WQQ2\|A0A084WQQ2_ANOSI Glutathione s-transferase E2 OS=Anopheles sinensis OX=74873 GN=ZHAS_00020698 PE=4 SV=1 |
| Query_10119 | tr\|A0A084VMZ1\|A0A084VMZ1_ANOSI Glutathione s-transferase E6 OS=Anopheles sinensis OX=74873 GN=ZHAS_00006695 PE=4 SV=1 |
| Query_10120 | tr\|A0A084WCJ1\|A0A084WCJ1_ANOSI AGAP011334-PA-like protein OS=Anopheles sinensis OX=74873 GN=ZHAS_00015997 PE=4 SV=1 |
| Query_10121 | tr\|A0A084WTI0\|A0A084WTI0_ANOSI AGAP004173-PA-like protein OS=Anopheles sinensis OX=74873 GN=ZHAS_00021753 PE=4 SV=1 |
| Query_10122 | tr\|A0A084WTJ9\|A0A084WTJ9_ANOSI AGAP004163-PB-like protein OS=Anopheles sinensis OX=74873 GN=ZHAS_00021774 PE=4 SV=1 |
| Query_10123 | sp\|Q93113\|GST1D_ANOGA Glutathione S-transferase 1, isoform D OS=Anopheles gambiae OX=7165 GN=GstD1 PE=1 SV=1 |
| Query_10124 | tr\|Q7QA79\|Q7QA79_ANOGA AGAP004383-PA OS=Anopheles gambiae OX=7165 GN=GSTD10 PE=4 SV=1 |
| Query_10125 | tr\|Q8MUS1\|Q8MUS1_ANOGA Glutathione S-transferase D11 OS=Anopheles gambiae OX=7165 GN=GSTd11 PE=2 SV=1 |
| Query_10126 | tr\|Q9GPL6\|Q9GPL6_ANOGA AGAP004380-PA OS=Anopheles gambiae OX=7165 GN=GSTD12 PE=2 SV=1 |
| Query_10127 | sp\|Q94999\|GSTT2_ANOGA Glutathione S-transferase 2 OS=Anopheles gambiae OX=7165 GN=GstD2 PE=3 SV=2 |
| Query_10128 | tr\|Q7PQ95\|Q7PQ95_ANOGA AGAP004382-PA OS=Anopheles gambiae OX=7165 GN=GSTD3 PE=3 SV=3 |
| Query_10129 | tr\|Q5TT03\|Q5TT03_ANOGA AGAP004381-PA OS=Anopheles gambiae OX=7165 GN=GSTD4 PE=3 SV=1 |
| Query_10130 | tr\|Q7QB59\|Q7QB59_ANOGA AGAP004173-PA OS=Anopheles gambiae OX=7165 GN=GSTD5 PE=4 SV=1 |
| Query_10131 | tr\|Q8MUS2\|Q8MUS2_ANOGA Glutathione S-transferase D6 (Fragment) OS=Anopheles gambiae OX=7165 GN=GSTd6 PE=2 SV=1 |
| Query_10132 | sp\|O76483\|GSTT7_ANOGA Glutathione S-transferase D7 OS=Anopheles gambiae OX=7165 GN=GstD7 PE=2 SV=1 |
| Query_10133 | tr\|Q5TTE5\|Q5TTE5_ANOGA AGAP004171-PA OS=Anopheles gambiae OX=7165 GN=GSTD8 PE=3 SV=2 |
| Query_10134 | tr\|Q86D84\|Q86D84_ANOGA AGAP004172-PA OS=Anopheles gambiae OX=7165 GN=GSTd9 PE=4 SV=1 |
| Query_10135 | tr\|Q9GPL9\|Q9GPL9_ANOGA AGAP009195-PA OS=Anopheles gambiae OX=7165 GN=GSTE1 PE=2 SV=1 |
| Query_10136 | tr\|Q7PVS6\|Q7PVS6_ANOGA AGAP009194-PA OS=Anopheles gambiae OX=7165 GN=GSTE2 PE=1 SV=3 |
| Query_10137 | tr\|Q8WQJ9\|Q8WQJ9_ANOGA AGAP009197-PA OS=Anopheles gambiae OX=7165 GN=GSTe3 PE=3 SV=1 |
| Query_10138 | tr\|Q8WQJ8\|Q8WQJ8_ANOGA AGAP009193-PA OS=Anopheles gambiae OX=7165 GN=GSTe4 PE=2 SV=1 |
| Query_10139 | tr\|Q8WQJ7\|Q8WQJ7_ANOGA AGAP009192-PA OS=Anopheles gambiae OX=7165 GN=GSTe5 PE=2 SV=1 |
| Query_10140 | tr\|A0NG89\|A0NG89_ANOGA AGAP009191-PA OS=Anopheles gambiae OX=7165 GN=GSTE6 PE=3 SV=1 |
| Query_10141 | tr\|Q7PVS4\|Q7PVS4_ANOGA AGAP009196-PA OS=Anopheles gambiae OX=7165 GN=GSTE7 PE=4 SV=3 |
| Query_10142 | tr\|Q8WQJ5\|Q8WQJ5_ANOGA AGAP009190-PA OS=Anopheles gambiae OX=7165 GN=GSTe8 PE=2 SV=1 |
| Query_10143 | tr\|Q8MUQ1\|Q8MUQ1_ANOGA AGAP000761-PA OS=Anopheles gambiae OX=7165 GN=gstT1 PE=2 SV=1 |
| Query_10144 | tr\|Q8MUQ2\|Q8MUQ2_ANOGA AGAP000888-PA OS=Anopheles gambiae OX=7165 GN=gstT2 PE=2 SV=1 |
| Query_10145 | tr\|A0A087ZVS9\|A0A087ZVS9_APIME Uncharacterized protein OS=Apis mellifera OX=7460 GN=GstT1 PE=3 SV=1 |
| Query_10146 | tr\|A0A088AFL5\|A0A088AFL5_APIME Uncharacterized protein OS=Apis mellifera OX=7460 GN=GstD1 PE=4 SV=1 |
| Query_10147 | tr\|Q6IVB7\|Q6IVB7_APILI Glutathione-S-transferase 1 (Fragment) OS=Apis mellifera ligustica OX=7469 GN=gst1 PE=2 SV=2 |
| Query_10148 | tr\|Q2I0J5\|Q2I0J5_BOMMO Glutathione S-transferase 3 OS=Bombyx mori OX=7091 GN=693114 PE=2 SV=1 |
| Query_10149 | tr\|H9IYE6\|H9IYE6_BOMMO Uncharacterized protein OS=Bombyx mori OX=7091 PE=3 SV=1 |
| Query_10150 | tr\|O61996\|O61996_BOMMO Glutathione S-transferase OS=Bombyx mori OX=7091 GN=692678 PE=1 SV=1 |
| Query_10151 | tr\|H9JAU4\|H9JAU4_BOMMO Uncharacterized protein OS=Bombyx mori OX=7091 PE=4 SV=1 |
| Query_10152 | tr\|H9JK84\|H9JK84_BOMMO Uncharacterized protein OS=Bombyx mori OX=7091 PE=3 SV=1 |
| Query_10153 | tr\|H9JKA0\|H9JKA0_BOMMO Uncharacterized protein OS=Bombyx mori OX=7091 PE=3 SV=1 |
| Query_10154 | tr\|H9JKP2\|H9JKP2_BOMMO Uncharacterized protein OS=Bombyx mori OX=7091 PE=3 SV=1 |
| Query_10155 | tr\|H9JKP3\|H9JKP3_BOMMO Uncharacterized protein OS=Bombyx mori OX=7091 PE=3 SV=1 |
| Query_10156 | BmoriGSTE7 |
| Query_10157 | tr\|B0LB16\|B0LB16_BOMMO Epsilon-class glutathione transferase OS=Bombyx mori OX=7091 GN=gste PE=2 SV=1 |
| Query_10158 | tr\|A0A077D6E7\|A0A077D6E7_CNAME Glutathione S-transferase delta 1 OS=Cnaphalocrocis medinalis OX=437488 PE=2 SV=1 |
| Query_10159 | tr\|A0A077D817\|A0A077D817_CNAME Glutathione S-transferase epsilon 1 OS=Cnaphalocrocis medinalis OX=437488 PE=2 SV=1 |
| Query_10160 | tr\|A0A0A7KNK1\|A0A0A7KNK1_CNAME Glutathione S-transferase delta 4 OS=Cnaphalocrocis medinalis OX=437488 PE=2 SV=1 |
| Query_10161 | tr\|A0A077D9Y8\|A0A077D9Y8_CNAME Glutathione S-transferase epsilon 2 OS=Cnaphalocrocis medinalis OX=437488 PE=2 SV=1 |
| Query_10162 | tr\|A0A0A7KL70\|A0A0A7KL70_CNAME Glutathione S-transferase epsilon 9 OS=Cnaphalocrocis medinalis OX=437488 PE=2 SV=1 |
| Query_10163 | tr\|A0A0A7KQB0\|A0A0A7KQB0_CNAME Glutathione S-transferase delta 3 OS=Cnaphalocrocis medinalis OX=437488 PE=2 SV=1 |
| Query_10164 | tr\|A0A077D602\|A0A077D602_CNAME Glutathione S-transferase OS=Cnaphalocrocis medinalis OX=437488 PE=2 SV=1 |
| Query_10165 | tr\|A0A0A7KLA5\|A0A0A7KLA5_CNAME Glutathione S-transferase epsilon 8 OS=Cnaphalocrocis medinalis OX=437488 PE=2 SV=1 |
| Query_10166 | tr\|A0A077D820\|A0A077D820_CNAME Glutathione S-transferase epsilon 6 (Fragment) OS=Cnaphalocrocis medinalis OX=437488 PE=2 SV=1 |
| Query_10167 | tr\|A0A077D5Z1\|A0A077D5Z1_CNAME Glutathione S-transferase epsilon 5 (Fragment) OS=Cnaphalocrocis medinalis OX=437488 PE=2 SV=1 |
| Query_10168 | tr\|A0A077D5Y7\|A0A077D5Y7_CNAME Glutathione S-transferase delta 2 OS=Cnaphalocrocis medinalis OX=437488 PE=2 SV=1 |
| Query_10169 | tr\|A0A077DB43\|A0A077DB43_CNAME Glutathione S-transferase epsilon 3 OS=Cnaphalocrocis medinalis OX=437488 PE=2 SV=1 |
| Query_10170 | tr\|A0A077D6F2\|A0A077D6F2_CNAME Glutathione S-transferase epsilon 4 OS=Cnaphalocrocis medinalis OX=437488 PE=2 SV=1 |
| Query_10171 | tr\|A0A0A7KLF5\|A0A0A7KLF5_CNAME Glutathione S-transferase epsilon 7 OS=Cnaphalocrocis medinalis OX=437488 PE=2 SV=1 |
| Query_10172 | tr\|B0VZJ3\|B0VZJ3_CULQU Glutathione S-transferase D2 OS=Culex quinquefasciatus OX=7176 GN=6031025 PE=3 SV=1 |
| Query_10173 | tr\|B0W6A9\|B0W6A9_CULQU Glutathione-s-transferase theta, gst OS=Culex quinquefasciatus OX=7176 GN=6033850 PE=4 SV=1 |
| Query_10174 | tr\|B0W6B2\|B0W6B2_CULQU Glutathione S-transferase 1-1 OS=Culex quinquefasciatus OX=7176 GN=6033854 PE=3 SV=1 |
| Query_10175 | tr\|B0W6C3\|B0W6C3_CULQU Glutathione S-transferase 1 OS=Culex quinquefasciatus OX=7176 GN=6033865 PE=4 SV=1 |
| Query_10176 | tr\|B0W6C4\|B0W6C4_CULQU Glutathione S-transferase 1 OS=Culex quinquefasciatus OX=7176 GN=6033866 PE=4 SV=1 |
| Query_10177 | tr\|B0W6C5\|B0W6C5_CULQU Glutathione S-transferase D7 OS=Culex quinquefasciatus OX=7176 GN=6033867 PE=3 SV=1 |
| Query_10178 | tr\|B0W6C7\|B0W6C7_CULQU Glutathione transferase I OS=Culex quinquefasciatus OX=7176 GN=6033869 PE=3 SV=1 |
| Query_10179 | tr\|B0W6C8\|B0W6C8_CULQU Glutathione S-transferase theta-2 OS=Culex quinquefasciatus OX=7176 GN=6033870 PE=3 SV=1 |
| Query_10180 | tr\|B0W6C9\|B0W6C9_CULQU Glutathione S-transferase OS=Culex quinquefasciatus OX=7176 GN=6033871 PE=4 SV=1 |
| Query_10181 | tr\|B0W6D0\|B0W6D0_CULQU Glutathione S-transferase OS=Culex quinquefasciatus OX=7176 GN=6033872 PE=3 SV=1 |
| Query_10182 | tr\|A0A1S4J718\|A0A1S4J718_CULQU GSTD2 protein OS=Culex quinquefasciatus OX=7176 PE=3 SV=1 |
| Query_10183 | tr\|B0W6D2\|B0W6D2_CULQU Glutathione S-transferase 1-1 OS=Culex quinquefasciatus OX=7176 GN=6033874 PE=3 SV=1 |
| Query_10184 | tr\|B0WQW9\|B0WQW9_CULQU Glutathione S-transferase 1 OS=Culex quinquefasciatus OX=7176 GN=6041912 PE=3 SV=1 |
| Query_10185 | tr\|B0WUG8\|B0WUG8_CULQU Glutathione S-transferase 1-5 OS=Culex quinquefasciatus OX=7176 GN=6043364 PE=3 SV=1 |
| Query_10186 | tr\|B0X3C7\|B0X3C7_CULQU Glutathione-s-transferase theta, gst OS=Culex quinquefasciatus OX=7176 GN=6047041 PE=3 SV=1 |
| Query_10187 | tr\|B0X3C8\|B0X3C8_CULQU Glutathione-s-transferase theta, gst OS=Culex quinquefasciatus OX=7176 GN=6047042 PE=3 SV=1 |
| Query_10188 | tr\|B0X3C9\|B0X3C9_CULQU Glutathione-s-transferase theta, gst OS=Culex quinquefasciatus OX=7176 GN=6047043 PE=3 SV=1 |
| Query_10189 | tr\|B0X3D0\|B0X3D0_CULQU Glutathione S-transferase theta-1 OS=Culex quinquefasciatus OX=7176 GN=6047044 PE=3 SV=1 |
| Query_10190 | tr\|B0XGJ6\|B0XGJ6_CULQU Glutathione-s-transferase theta, gst OS=Culex quinquefasciatus OX=7176 GN=6052508 PE=3 SV=1 |
| Query_10191 | tr\|B0XGJ7\|B0XGJ7_CULQU Glutathione-s-transferase theta, gst OS=Culex quinquefasciatus OX=7176 GN=6052509 PE=4 SV=1 |
| Query_10192 | tr\|B0XGJ8\|B0XGJ8_CULQU Glutathione S-transferase 1-1 OS=Culex quinquefasciatus OX=7176 GN=6052510 PE=3 SV=1 |
| Query_10193 | tr\|B0XGJ9\|B0XGJ9_CULQU Glutathione S-transferase E2 OS=Culex quinquefasciatus OX=7176 GN=6052511 PE=4 SV=1 |
| Query_10194 | tr\|B0XGK0\|B0XGK0_CULQU Glutathione-s-transferase theta, gst OS=Culex quinquefasciatus OX=7176 GN=6052512 PE=3 SV=1 |
| Query_10195 | tr\|B0XGK1\|B0XGK1_CULQU Glutathione S-transferase 1-1 OS=Culex quinquefasciatus OX=7176 GN=6052513 PE=4 SV=1 |
| Query_10196 | tr\|B0XGK2\|B0XGK2_CULQU Glutathione-s-transferase theta, gst OS=Culex quinquefasciatus OX=7176 GN=6052515 PE=4 SV=1 |
| Query_10197 | tr\|B0XGK3\|B0XGK3_CULQU Glutathione-s-transferase theta, gst OS=Culex quinquefasciatus OX=7176 GN=6052516 PE=4 SV=1 |
| Query_10198 | tr\|B0XGK4\|B0XGK4_CULQU Glutathione-s-transferase theta OS=Culex quinquefasciatus OX=7176 GN=6052517 PE=4 SV=1 |
| Query_10199 | tr\|B0XJU4\|B0XJU4_CULQU Glutathione transferase AtGST OS=Culex quinquefasciatus OX=7176 GN=6053908 PE=4 SV=1 |
| Query_10200 | tr\|B0XLC5\|B0XLC5_CULQU Glutathione transferase AtGST OS=Culex quinquefasciatus OX=7176 GN=6054553 PE=3 SV=1 |
| Query_10201 | DPOGS200212-PA |
| Query_10202 | DPOGS202622-PA |
| Query_10203 | DPOGS204831-PA |
| Query_10204 | DPOGS207576-PA |
| Query_10205 | DPOGS207703-PA |
| Query_10206 | DPOGS208312-PA |
| Query_10207 | DPOGS209578-PA |
| Query_10208 | DPOGS210477-PA |
| Query_10209 | DPOGS210488-PA |
| Query_10210 | DPOGS210526-PA |
| Query_10211 | sp\|P20432\|GSTD1_DROME Glutathione S-transferase D1 OS=Drosophila melanogaster GN=GstD1 PE=1 SV=1 |
| Query_10212 | tr\|Q9VGA1\|Q9VGA1_DROME Glutathione S transferase D10, isoform A OS=Drosophila melanogaster GN=GstD10 PE=1 SV=1 |
| Query_10213 | tr\|B7Z0S9\|B7Z0S9_DROME Glutathione S transferase D11, isoform B OS=Drosophila melanogaster GN=GstD11 PE=4 SV=1 |
| Query_10214 | tr\|Q8SXQ9\|Q8SXQ9_DROME Glutathione S transferase D11, isoform A OS=Drosophila melanogaster GN=GstD11 PE=2 SV=1 |
| Query_10215 | sp\|Q9VG98\|GSTD2_DROME Glutathione S-transferase D2 OS=Drosophila melanogaster GN=GstD2 PE=1 SV=1 |
| Query_10216 | sp\|Q9VG97\|GSTD3_DROME Inactive glutathione S-transferase D3 OS=Drosophila melanogaster GN=GstD3 PE=2 SV=1 |
| Query_10217 | sp\|Q9VG96\|GSTD4_DROME Glutathione S-transferase D4 OS=Drosophila melanogaster GN=GstD4 PE=1 SV=1 |
| Query_10218 | sp\|Q9VG95\|GSTD5_DROME Glutathione S-transferase D5 OS=Drosophila melanogaster GN=GstD5 PE=1 SV=2 |
| Query_10219 | sp\|Q9VG94\|GSTD6_DROME Glutathione S-transferase D6 OS=Drosophila melanogaster GN=GstD6 PE=1 SV=1 |
| Query_10220 | sp\|Q9VG93\|GSTD7_DROME Glutathione S-transferase D7 OS=Drosophila melanogaster GN=GstD7 PE=1 SV=1 |
| Query_10221 | tr\|Q9VG92\|Q9VG92_DROME Glutathione S transferase D8 OS=Drosophila melanogaster GN=GstD8 PE=2 SV=1 |
| Query_10222 | tr\|Q9VGA0\|Q9VGA0_DROME Glutathione S transferase D9, isoform A OS=Drosophila melanogaster GN=GstD9 PE=1 SV=1 |
| Query_10223 | tr\|Q7KK90\|Q7KK90_DROME GH14654p OS=Drosophila melanogaster GN=GstE1 PE=1 SV=1 |
| Query_10224 | tr\|Q4V6J1\|Q4V6J1_DROME Glutathione S transferase E10, isoform A OS=Drosophila melanogaster GN=GstE10 PE=2 SV=1 |
| Query_10225 | tr\|Q7JVZ8\|Q7JVZ8_DROME Glutathione S transferase E11, isoform A OS=Drosophila melanogaster GN=GstE11 PE=1 SV=1 |
| Query_10226 | tr\|Q9XYZ9\|Q9XYZ9_DROME Glutathione S transferase E12, isoform A OS=Drosophila melanogaster GN=GstE12 PE=1 SV=1 |
| Query_10227 | tr\|Q7JVI6\|Q7JVI6_DROME Glutathione S transferase E13, isoform A OS=Drosophila melanogaster GN=GstE13 PE=1 SV=1 |
| Query_10228 | sp\|Q7JYX0\|GSTEE_DROME Glutathione S-transferase E14 OS=Drosophila melanogaster GN=GstE14 PE=1 SV=1 |
| Query_10229 | tr\|Q7JYZ9\|Q7JYZ9_DROME Glutathione S transferase E2 OS=Drosophila melanogaster GN=GstE2 PE=2 SV=1 |
| Query_10230 | tr\|A1ZB68\|A1ZB68_DROME FI01423p OS=Drosophila melanogaster GN=GstE3 PE=1 SV=1 |
| Query_10231 | tr\|A1ZB69\|A1ZB69_DROME Glutathione S transferase E4 OS=Drosophila melanogaster GN=GstE4 PE=3 SV=1 |
| Query_10232 | tr\|A1ZB70\|A1ZB70_DROME Glutathione S transferase E5 OS=Drosophila melanogaster GN=GstE5 PE=3 SV=1 |
| Query_10233 | tr\|A1ZB71\|A1ZB71_DROME Glutathione S transferase E6 OS=Drosophila melanogaster GN=GstE6 PE=1 SV=1 |
| Query_10234 | tr\|A1ZB72\|A1ZB72_DROME Glutathione S transferase E7 OS=Drosophila melanogaster GN=GstE7 PE=1 SV=1 |
| Query_10235 | tr\|A1ZB73\|A1ZB73_DROME Glutathione S transferase E8, isoform A OS=Drosophila melanogaster GN=GstE8 PE=3 SV=1 |
| Query_10236 | tr\|Q7K8X7\|Q7K8X7_DROME Glutathione S transferase E9 OS=Drosophila melanogaster GN=GstE9 PE=1 SV=1 |
| Query_10237 | tr\|Q7K0B6\|Q7K0B6_DROME Glutathione S transferase T1 OS=Drosophila melanogaster OX=7227 GN=GstT1 PE=1 SV=1 |
| Query_10238 | tr\|A1Z7X7\|A1Z7X7_DROME Glutathione S transferase T2 OS=Drosophila melanogaster OX=7227 GN=GstT2 PE=1 SV=2 |
| Query_10239 | tr\|Q9VRA4\|Q9VRA4_DROME Glutathione S transferase T3, isoform A OS=Drosophila melanogaster OX=7227 GN=GstT3 PE=3 SV=2 |
| Query_10240 | tr\|E1JJS1\|E1JJS1_DROME Glutathione S transferase T3, isoform B OS=Drosophila melanogaster OX=7227 GN=GstT3 PE=3 SV=1 |
| Query_10241 | tr\|Q8MRM0\|Q8MRM0_DROME GH16740p OS=Drosophila melanogaster OX=7227 GN=GstT4 PE=1 SV=1 |
| Query_10242 | tr\|A0A291ARU4\|A0A291ARU4_HELAM Glutathione S-transferase OS=Helicoverpa armigera OX=29058 GN=GST8 PE=2 SV=1 |
| Query_10243 | tr\|A0A2W1BRB0\|A0A2W1BRB0_HELAM Uncharacterized protein OS=Helicoverpa armigera OX=29058 GN=HaOG200226 PE=3 SV=1 |
| Query_10244 | tr\|A0A2W1BSA5\|A0A2W1BSA5_HELAM Uncharacterized protein OS=Helicoverpa armigera OX=29058 GN=HaOG200219 PE=3 SV=1 |
| Query_10245 | tr\|A0A2W1BUA9\|A0A2W1BUA9_HELAM Uncharacterized protein OS=Helicoverpa armigera OX=29058 GN=HaOG200217 PE=3 SV=1 |
| Query_10246 | tr\|A0A2W1BZ02\|A0A2W1BZ02_HELAM Uncharacterized protein OS=Helicoverpa armigera OX=29058 GN=HaOG200220 PE=3 SV=1 |
| Query_10247 | tr\|A0MSN0\|A0MSN0_HELAM Glutathione S-transferase OS=Helicoverpa armigera OX=29058 PE=2 SV=1 |
| Query_10248 | tr\|B6A8L4\|B6A8L4_HELAM Glutathione S-transferase (Fragment) OS=Helicoverpa armigera OX=29058 PE=2 SV=1 |
| Query_10249 | tr\|C8YL89\|C8YL89_HELAM Glutathione S-transferase 16 OS=Helicoverpa armigera OX=29058 GN=GST16 PE=2 SV=1 |
| Query_10250 | tr\|D7NI45\|D7NI45_HELAM Glutathione S-transferase OS=Helicoverpa armigera OX=29058 GN=GST6 PE=2 SV=1 |
| Query_10251 | AIB07715.1 glutathione S-transferase GSTD1 [Helicoverpa armigera armigera] |
| Query_10252 | AIB07714.1 glutathione S-transferase GSTD3 [Helicoverpa armigera armigera] |
| Query_10253 | AIB07716.1 glutathione S-transferase GSTD4 [Helicoverpa armigera armigera] |
| Query_10254 | AIB07717.1 glutathione S-transferase GSTD5, partial [Helicoverpa armigera armigera] |
| Query_10255 | XP_021189521.1 glutathione S-transferase D7-like isoform X1 [Helicoverpa armigera] |
| Query_10256 | Helicoverpa_armigera_GSTE14-like_isoform_X2 |
| Query_10257 | tr\|Q7Z0Q7\|Q7Z0Q7_HELAM Glutathione S-transferase (Fragment) OS=Helicoverpa armigera OX=29058 PE=2 SV=1 |
| Query_10258 | tr\|Q7Z0Q8\|Q7Z0Q8_HELAM Glutathione S-transferase (Fragment) OS=Helicoverpa armigera OX=29058 PE=2 SV=1 |
| Query_10259 | tr\|Q7Z0Q9\|Q7Z0Q9_HELAM Glutathione S-transferase (Fragment) OS=Helicoverpa armigera OX=29058 PE=2 SV=1 |
| Query_10260 | tr\|A0A2A4JDE9\|A0A2A4JDE9_HELVI Uncharacterized protein OS=Heliothis virescens OX=7102 GN=B5V51_3669 PE=3 SV=1 |
| Query_10261 | tr\|A0A2A4IXQ2\|A0A2A4IXQ2_HELVI Uncharacterized protein OS=Heliothis virescens OX=7102 GN=B5V51_11472 PE=3 SV=1 |
| Query_10262 | tr\|A0A2A4JI76\|A0A2A4JI76_HELVI Uncharacterized protein OS=Heliothis virescens OX=7102 GN=B5V51_1534 PE=3 SV=1 |
| Query_10263 | tr\|A0A2A4JDN1\|A0A2A4JDN1_HELVI Uncharacterized protein OS=Heliothis virescens OX=7102 GN=B5V51_3670 PE=3 SV=1 |
| Query_10264 | tr\|A0A2A4JU40\|A0A2A4JU40_HELVI Uncharacterized protein OS=Heliothis virescens OX=7102 GN=B5V51_11931 PE=3 SV=1 |
| Query_10265 | tr\|A0A2A4IV17\|A0A2A4IV17_HELVI Uncharacterized protein OS=Heliothis virescens OX=7102 GN=B5V51_12180 PE=3 SV=1 |
| Query_10266 | tr\|A0A2A4IXK8\|A0A2A4IXK8_HELVI Uncharacterized protein OS=Heliothis virescens OX=7102 GN=B5V51_10494 PE=3 SV=1 |
| Query_10267 | tr\|A0A2A4JCV1\|A0A2A4JCV1_HELVI Uncharacterized protein OS=Heliothis virescens OX=7102 GN=B5V51_3945 PE=3 SV=1 |
| Query_10268 | tr\|A0A2A4JV70\|A0A2A4JV70_HELVI Uncharacterized protein OS=Heliothis virescens OX=7102 GN=B5V51_11943 PE=3 SV=1 |
| Query_10269 | tr\|A0A2A4IY34\|A0A2A4IY34_HELVI Uncharacterized protein (Fragment) OS=Heliothis virescens OX=7102 GN=B5V51_11290 PE=4 SV=1 |
| Query_10270 | tr\|A0A2A4JKT9\|A0A2A4JKT9_HELVI Uncharacterized protein OS=Heliothis virescens OX=7102 GN=B5V51_845 PE=4 SV=1 |
| Query_10271 | NP_001165913.1 glutathione S-transferase D1 [Nasonia vitripennis] |
| Query_10272 | NP_001165914.1 glutathione S-transferase D3 [Nasonia vitripennis] |
| Query_10273 | NP_001165915.1 glutathione S-transferase D5 [Nasonia vitripennis] |
| Query_10274 | NP_001165925.1 glutathione S-transferase T2 [Nasonia vitripennis] |
| Query_10275 | NP_001165926.1 glutathione S-transferase T1 [Nasonia vitripennis] |
| Query_10276 | NP_001165927.1 glutathione S-transferase T3 [Nasonia vitripennis] |
| Query_10277 | XP_001600187.1 PREDICTED: glutathione S-transferase D7 [Nasonia vitripennis] |
| Query_10278 | tr\|A0A0L7KM86\|A0A0L7KM86_9NEOP Glutathione S-transferase OS=Operophtera brumata OX=104452 GN=OBRU01_24524 PE=3 SV=1 |
| Query_10279 | tr\|A0A0L7LFQ3\|A0A0L7LFQ3_9NEOP Glutathione S-transferase epsilon (Fragment) OS=Operophtera brumata OX=104452 GN=OBRU01_09353 PE=4 SV=1 |
| Query_10280 | tr\|A0A0L7LME0\|A0A0L7LME0_9NEOP Glutathione S-transferase 1-6 (Fragment) OS=Operophtera brumata OX=104452 GN=OBRU01_05580 PE=3 SV=1 |
| Query_10281 | tr\|A0A0L7LPT4\|A0A0L7LPT4_9NEOP Glutathione S-transferase OS=Operophtera brumata OX=104452 GN=OBRU01_04185 PE=3 SV=1 |
| Query_10282 | tr\|A0A0L7LTB4\|A0A0L7LTB4_9NEOP Glutathione S-transferase OS=Operophtera brumata OX=104452 GN=OBRU01_01600 PE=3 SV=1 |
| Query_10283 | tr\|A0A0L7LTL7\|A0A0L7LTL7_9NEOP Glutathione S-transferase epsilon 11 OS=Operophtera brumata OX=104452 GN=OBRU01_01599 PE=3 SV=1 |
| Query_10284 | tr\|A0A0L7LI53\|A0A0L7LI53_9NEOP Glutathione S-transferase epsilon 6 (Fragment) OS=Operophtera brumata OX=104452 GN=OBRU01_07826 PE=4 SV=1 |
| Query_10285 | tr\|A0A0L7KLM3\|A0A0L7KLM3_9NEOP Glutathione S-transferase 10 OS=Operophtera brumata OX=104452 GN=OBRU01_24503 PE=4 SV=1 |
| Query_10286 | tr\|A0A0L7L1M8\|A0A0L7L1M8_9NEOP Glutathione S-transferase epsilon 3 OS=Operophtera brumata OX=104452 GN=OBRU01_16992 PE=4 SV=1 |
| Query_10287 | tr\|A0A0L7LM54\|A0A0L7LM54_9NEOP Glutathione S-transferase 1 OS=Operophtera brumata OX=104452 GN=OBRU01_05579 PE=4 SV=1 |
| Query_10288 | tr\|A0A194QXR6\|A0A194QXR6_PAPMA Glutathione S-transferase D7 OS=Papilio machaon OX=76193 GN=RR48_13110 PE=4 SV=1 |
| Query_10289 | tr\|A0A194R5D8\|A0A194R5D8_PAPMA Glutathione S-transferase 1 OS=Papilio machaon OX=76193 GN=RR48_10522 PE=4 SV=1 |
| Query_10290 | tr\|A0A0N1PJW2\|A0A0N1PJW2_PAPMA Glutathione S-transferase theta-1 OS=Papilio machaon OX=76193 GN=RR48_04090 PE=3 SV=1 |
| Query_10291 | tr\|A0A194QX62\|A0A194QX62_PAPMA Glutathione S-transferase 1, isoform D OS=Papilio machaon OX=76193 GN=RR48_13184 PE=3 SV=1 |
| Query_10292 | tr\|A0A194REA6\|A0A194REA6_PAPMA Glutathione S-transferase 1-1 OS=Papilio machaon OX=76193 GN=RR48_07011 PE=3 SV=1 |
| Query_10293 | XP_014368559.1 PREDICTED: glutathione S-transferase E14-like [Papilio machaon] |
| Query_10294 | tr\|I4DRD9\|I4DRD9_PAPPL Glutathione S transferase D8 OS=Papilio polytes OX=76194 PE=2 SV=1 |
| Query_10295 | tr\|I4DRZ4\|I4DRZ4_PAPPL Glutathione S transferase E8 (Fragment) OS=Papilio polytes OX=76194 PE=2 SV=1 |
| Query_10296 | XP_013145092.1 PREDICTED: glutathione S-transferase theta-1-like [Papilio polytes] |
| Query_10297 | XP_013142369.1 PREDICTED: glutathione S-transferase 1, isoform D-like [Papilio polytes] |
| Query_10298 | XP_013142272.1 PREDICTED: glutathione S-transferase D7-like isoform X2 [Papilio polytes] |
| Query_10299 | Papilio_polytes_GST1-1-like_isoform_X1 |
| Query_10300 | tr\|A0A194PX26\|A0A194PX26_PAPXU Glutathione S-transferase theta-1 OS=Papilio xuthus OX=66420 GN=RR46_11408 PE=3 SV=1 |
| Query_10301 | tr\|Q4R1I6\|Q4R1I6_PAPXU Glutathione-S-transferase OS=Papilio xuthus OX=66420 GN=GST-pxcs1 PE=3 SV=1 |
| Query_10302 | tr\|I4DKS5\|I4DKS5_PAPXU Glutathionetransferase OS=Papilio xuthus OX=66420 PE=2 SV=1 |
| Query_10303 | tr\|A0A194QHF9\|A0A194QHF9_PAPXU Glutathione S-transferase 1-1 OS=Papilio xuthus OX=66420 GN=RR46_01739 PE=3 SV=1 |
| Query_10304 | tr\|I4DNY4\|I4DNY4_PAPXU Glutathionetransferase OS=Papilio xuthus OX=66420 PE=2 SV=1 |
| Query_10305 | tr\|A0A194QG68\|A0A194QG68_PAPXU Glutathione S-transferase D7 OS=Papilio xuthus OX=66420 GN=RR46_08238 PE=3 SV=1 |
| Query_10306 | tr\|I4DQ12\|I4DQ12_PAPXU Glutathione S transferase E6 (Fragment) OS=Papilio xuthus OX=66420 PE=2 SV=1 |
| Query_10307 | tr\|E0VCV3\|E0VCV3_PEDHC GSTD1-5 protein, putative OS=Pediculus humanus subsp. corporis OX=121224 GN=8238066 PE=4 SV=1 |
| Query_10308 | tr\|E0VGM2\|E0VGM2_PEDHC GSTD1-5 protein, putative OS=Pediculus humanus subsp. corporis OX=121224 GN=8240051 PE=3 SV=1 |
| Query_10309 | tr\|E0VGM3\|E0VGM3_PEDHC GSTD1-5 protein, putative OS=Pediculus humanus subsp. corporis OX=121224 GN=8240052 PE=3 SV=1 |
| Query_10310 | tr\|E0VUR9\|E0VUR9_PEDHC GSTD1-5 protein, putative OS=Pediculus humanus subsp. corporis OX=121224 GN=8230516 PE=3 SV=1 |
| Query_10311 | tr\|A0A1P8L0T5\|A0A1P8L0T5_PIERA Glutathione S-transferase theta 1 OS=Pieris rapae OX=64459 PE=2 SV=1 |
| Query_10312 | tr\|A0A1P8L0U3\|A0A1P8L0U3_PIERA Glutathione S-transferase epsilon 1 OS=Pieris rapae OX=64459 PE=2 SV=1 |
| Query_10313 | tr\|A0A1P8L0T0\|A0A1P8L0T0_PIERA Glutathione S-transferase epsilon 2 OS=Pieris rapae OX=64459 PE=2 SV=1 |
| Query_10314 | tr\|A0A1P8L0T7\|A0A1P8L0T7_PIERA Glutathione S-transferase OS=Pieris rapae OX=64459 PE=2 SV=1 |
| Query_10315 | tr\|A0A1P8L0S7\|A0A1P8L0S7_PIERA Glutathione S-transferase delta 2 OS=Pieris rapae OX=64459 PE=2 SV=1 |
| Query_10316 | tr\|A0A1P8L0S9\|A0A1P8L0S9_PIERA Glutathione S-transferase delta 1 OS=Pieris rapae OX=64459 PE=2 SV=1 |
| Query_10317 | tr\|A0A1P8L0S2\|A0A1P8L0S2_PIERA Glutathione S-transferase epsilon 3 OS=Pieris rapae OX=64459 PE=2 SV=1 |
| Query_10318 | XP_022126447.1 glutathione S-transferase E14-like [Pieris rapae] |
| Query_10319 | tr\|O77409\|O77409_PLUXY Glutathione S-transferase isozyme 3 OS=Plutella xylostella OX=51655 GN=GST3 PE=2 SV=1 |
| Query_10320 | tr\|Q2ABX5\|Q2ABX5_PLUXY Glutathione S-Transferase-Epsilon7 OS=Plutella xylostella OX=51655 GN=GST3 PE=2 SV=1 |
| Query_10321 | tr\|X5D044\|X5D044_PLUXY Glutathione S-transferase (Fragment) OS=Plutella xylostella OX=51655 PE=2 SV=1 |
| Query_10322 | tr\|X5CJS2\|X5CJS2_PLUXY Glutathione S-transferase OS=Plutella xylostella OX=51655 PE=2 SV=1 |
| Query_10323 | tr\|X5CYF6\|X5CYF6_PLUXY Glutathione S-transferase OS=Plutella xylostella OX=51655 PE=2 SV=1 |
| Query_10324 | tr\|X5CCB3\|X5CCB3_PLUXY Glutathione S-transferase OS=Plutella xylostella OX=51655 PE=2 SV=1 |
| Query_10325 | tr\|X5CYG4\|X5CYG4_PLUXY Glutathione S-transferase OS=Plutella xylostella OX=51655 PE=2 SV=1 |
| Query_10326 | tr\|X5CJR9\|X5CJR9_PLUXY Glutathione S-transferase OS=Plutella xylostella OX=51655 PE=2 SV=1 |
| Query_10327 | tr\|A0A1L8D6F5\|A0A1L8D6F5_PLUXY Glutathione S-Transferase-Epsilon6 OS=Plutella xylostella OX=51655 PE=2 SV=1 |
| Query_10328 | tr\|X5CHJ9\|X5CHJ9_PLUXY Glutathione S-transferase OS=Plutella xylostella OX=51655 PE=2 SV=1 |
| Query_10329 | tr\|D7URW9\|D7URW9_PLUXY Glutathione S-transferase delta OS=Plutella xylostella OX=51655 GN=PxGSTd PE=2 SV=1 |
| Query_10330 | tr\|X5CCB0\|X5CCB0_PLUXY Glutathione S-transferase OS=Plutella xylostella OX=51655 PE=2 SV=1 |
| Query_10331 | tr\|X5CJR6\|X5CJR6_PLUXY Glutathione S-transferase OS=Plutella xylostella OX=51655 PE=2 SV=1 |
| Query_10332 | tr\|X5D048\|X5D048_PLUXY Glutathione S-transferase OS=Plutella xylostella OX=51655 PE=2 SV=1 |
| Query_10333 | tr\|D2I931\|D2I931_SPOLT Glutathione S-transferase epsilon 3 OS=Spodoptera litura OX=69820 PE=2 SV=1 |
| Query_10334 | tr\|D2I930\|D2I930_SPOLT Glutathione S-transferase epsilon 2 OS=Spodoptera litura OX=69820 PE=2 SV=1 |
| Query_10335 | tr\|A0A075X2X0\|A0A075X2X0_SPOLT Glutathione S-transferase theta 1 OS=Spodoptera litura OX=69820 PE=2 SV=1 |
| Query_10336 | tr\|A0A075X3S8\|A0A075X3S8_SPOLT Glutathione S-transferase delta 4 OS=Spodoptera litura OX=69820 PE=2 SV=1 |
| Query_10337 | tr\|A0A075X2I2\|A0A075X2I2_SPOLT Glutathione S-transferase epsilon 4 OS=Spodoptera litura OX=69820 PE=2 SV=1 |
| Query_10338 | tr\|A0A075X2W2\|A0A075X2W2_SPOLT Glutathione S-transferase epsilon 15 OS=Spodoptera litura OX=69820 PE=2 SV=1 |
| Query_10339 | tr\|A0A075X8X2\|A0A075X8X2_SPOLT Glutathione S-transferase epsilon 6 OS=Spodoptera litura OX=69820 PE=2 SV=1 |
| Query_10340 | tr\|A0A075X8Y6\|A0A075X8Y6_SPOLT Glutathione S-transferase delta 3 OS=Spodoptera litura OX=69820 PE=2 SV=1 |
| Query_10341 | tr\|A0A075X244\|A0A075X244_SPOLT Glutathione S-transferase epsilon 8 OS=Spodoptera litura OX=69820 PE=2 SV=1 |
| Query_10342 | tr\|A0A075X8X7\|A0A075X8X7_SPOLT Glutathione S-transferase epsilon 11 OS=Spodoptera litura OX=69820 PE=2 SV=1 |
| Query_10343 | tr\|Q1EGY7\|Q1EGY7_SPOLT Gst1 OS=Spodoptera litura OX=69820 GN=gst1 PE=2 SV=1 |
| Query_10344 | tr\|A0A075X250\|A0A075X250_SPOLT Glutathione S-transferase epsilon 13 (Fragment) OS=Spodoptera litura OX=69820 PE=2 SV=1 |
| Query_10345 | tr\|A0A075X2W6\|A0A075X2W6_SPOLT Glutathione S-transferase delta 2 (Fragment) OS=Spodoptera litura OX=69820 PE=2 SV=1 |
| Query_10346 | tr\|A0A075X2V8\|A0A075X2V8_SPOLT Glutathione S-transferase epsilon 10 (Fragment) OS=Spodoptera litura OX=69820 PE=2 SV=1 |
| Query_10347 | tr\|A0A075X2J6\|A0A075X2J6_SPOLT Glutathione S-transferase delta 1 (Fragment) OS=Spodoptera litura OX=69820 PE=2 SV=1 |
| Query_10348 | tr\|A0A075X2I7\|A0A075X2I7_SPOLT Glutathione S-transferase epsilon 9 OS=Spodoptera litura OX=69820 PE=2 SV=1 |
| Query_10349 | tr\|A0A075X3R3\|A0A075X3R3_SPOLT Glutathione S-transferase epsilon 7 (Fragment) OS=Spodoptera litura OX=69820 PE=2 SV=1 |
| Query_10350 | tr\|A0A075X2V3\|A0A075X2V3_SPOLT Glutathione S-transferase epsilon 5 OS=Spodoptera litura OX=69820 PE=2 SV=1 |
| Query_10351 | tr\|A0A075X3R8\|A0A075X3R8_SPOLT Glutathione S-transferase epsilon 12 OS=Spodoptera litura OX=69820 PE=2 SV=1 |
| Query_10352 | tr\|A0A075X2J2\|A0A075X2J2_SPOLT Glutathione S-transferase epsilon 14 OS=Spodoptera litura OX=69820 PE=2 SV=1 |
| Query_10353 | tr\|A0A077D0A2\|A0A077D0A2_SPOLT Glutathione S-transferase epsilon 3 (Fragment) OS=Spodoptera litura OX=69820 PE=2 SV=1 |
| Query_10354 | XP_022837684.1 glutathione S-transferase E14-like isoform X1 [Spodoptera litura] |
| Query_10355 | TC003103_001 peptide: TC003103_001 pep:protein_coding |
| Query_10356 | TC003104_001 peptide: TC003104_001 pep:protein_coding |
| Query_10357 | TC003345_001 peptide: TC003345_001 pep:protein_coding |
| Query_10358 | TC003347_001 peptide: TC003347_001 pep:protein_coding |
| Query_10359 | TC003348_001 peptide: TC003348_001 pep:protein_coding |
| Query_10360 | TC004442_001 peptide: TC004442_001 pep:protein_coding |
| Query_10361 | TC004443_001 peptide: TC004443_001 pep:protein_coding |
| Query_10362 | TC004444_001 peptide: TC004444_001 pep:protein_coding |
| Query_10363 | TC004447_001 peptide: TC004447_001 pep:protein_coding |
| Query_10364 | TC004448_001 peptide: TC004448_001 pep:protein_coding |
| Query_10365 | TC004449_001 peptide: TC004449_001 pep:protein_coding |
| Query_10366 | TC004450_001 peptide: TC004450_001 pep:protein_coding |
| Query_10367 | TC004940_001 peptide: TC004940_001 pep:protein_coding |
| Query_10368 | TC004941_001 peptide: TC004941_001 pep:protein_coding |
| Query_10369 | TC004942_001 peptide: TC004942_001 pep:protein_coding |
| Query_10370 | TC006215_001 peptide: TC006215_001 pep:protein_coding |
| Query_10371 | TC009482_001 peptide: TC009482_001 pep:protein_coding |
| Query_10372 | tr\|A0A2W1BRE1\|A0A2W1BRE1_HELAM Uncharacterized protein OS=Helicoverpa armigera OX=29058 GN=HaOG200227 PE=3 SV=1 |

**Movie Legends**

**Movie 1. Trajectory of MD simulations of DmNobo[WT]_EST-GSH or DmNobo[Asp113Ala]_EST-GSH.**

The Cα atoms of chain A of DmNobo[WT]_EST-GSH (blue) or DmNobo[Asp113Ala]_EST-GSH (white) in each frame were superimposed for 100 ns with those of DmNobo_EST-GSH at the initial state. The main chain of chain A of the protein is shown by a ribbon. Asp113 in DmNobo[WT]_EST-GSH, Ala113 in DmNobo[Asp113Ala]_EST-GSH, GSH, and EST are represented by sticks. Carbon atoms of EST in DmNobo[WT]_EST-GSH and those in DmNobo[Asp113Ala]_EST-GSH are colored in blue and magenta, respectively. The colors are the same as indicated in Fig. 4*D*.

**Movie 2. An enlarged view of the trajectory of EST in MD simulations for DmNobo[WT]_EST-GSH or DmNobo[Asp113Ala]_EST-GSH.**

An enlarged view of the trajectory of EST in MD simulations of DmNobo[WT]_EST-GSH or DmNobo[Asp113Ala]_EST-GSH (Movie 1)
